# The Fossilised Birth-Death Model is Identifiable

**DOI:** 10.1101/2024.02.08.579547

**Authors:** Kate Truman, Timothy G Vaughan, Alex Gavryushkin, Alexandra “Sasha” Gavryushkina

**Affiliations:** Biological Data Science Laboratory, School of Mathematics and Statistics, University of Canterbury, Christchurch, New Zealand; Biomathematics Research Centre, University of Canterbury, Christchurch, New Zealand; Department of Biosystems Science and Engineering, ETH Zurich, Basel 4056, Switzerland; Swiss Institute of Bioinformatics, Lausanne 1015, Switzerland

**Keywords:** Birth-death sampling models, Phylogenetics, Fossils, Speciation, Extinction

## Abstract

Time-dependent birth-death sampling models have been used in numerous studies for inferring past evolutionary dynamics in different areas, e.g. speciation and extinction rates in macroevolutionary studies, or effective reproductive number in epidemiological studies. These models are branching processes where lineages can bifurcate, die, or be sampled with time-dependent birth, death, and sampling rates, generating phylogenetic trees. It has been shown that in some subclasses of such models, different sets of rates can result in the same distributions of reconstructed phylogenetic trees, and therefore the rates become unidentifiable from the trees regardless of their size. Here we show that widely used time-dependent fossilised birth-death (FBD) models are identifiable. This subclass of models makes more realistic assumptions about the fossilisation process and certain infectious disease transmission processes than the unidentifiable birth-death sampling models. Namely, FBD models assume that sampled lineages stay in the process rather than being immediately removed upon sampling. Identifiability of the time-dependent FBD model justifies using statistical methods that implement this model to infer the underlying temporal diversification or epidemiological dynamics from phylogenetic trees or directly from molecular or other comparative data. We further show that the time-dependent fossilised-birth-death model with an extra parameter, the removal after sampling probability, is unidentifiable. This implies that in scenarios where we do not know how sampling affects lineages we are unable to infer this extra parameter together with birth, death, and sampling rates solely from trees.

## 1 Introduction

Birth-death models have been widely used to model different processes that are described by phylogenetic trees, from evolutionary histories of species (Raup et al., 1973; Nee et al., 1995; Nee, 2006; Heath et al., 2014; Silvestro et al., 2014; Morlon et al., 2011) to infectious disease transmission histories (Stadler et al., 2012, 2013). A birth-death process describes how lineages bifurcate and die with corresponding birth and death rates, and results in a phylogenetic tree (Feller, 1939; Kendall, 1948). Given molecular or other data of sampled lineages one can reconstruct a phylogenetic tree, and given the tree infer the rates, or infer trees and rates in a joint analysis (Yang and Rannala, 1997; Höhna et al., 2016; Bouckaert et al., 2019). The birth and death rates or their transformations are then interpreted as parameters of the underlying processes, e.g. speciation and extinction rates in macroevolutionary studies or effective reproductive number in epidemiological applications.

An important aspect of modelling the process that produces phylogenetic trees is the fact that such trees are reconstructed from a sample of extant species and do not represent all lineages that are involved in the evolutionary process. Extinct or unsampled lineages remain unobserved. To model present sampling, additionally to the birth and death rates we have a sampling fraction parameter (Stadler, 2009). Furthermore, we consider reconstructed trees that represent the observed part of the evolutionary history (Nee et al., 1995), that is, a phylogeny that relates sampled extant species only.

Temporal changes in the rates of species diversification or infectious disease transmission have been accounted for by time-dependent birth and death rates (Kendall, 1948; Stadler et al., 2013). The probability density functions for reconstructed trees, in this case, were derived for the simple case of piecewise constant rates (Stadler, 2011) or more general case with arbitrary time-dependent rates (Morlon et al., 2011), which enabled efficient inference of the speciation or transmission temporal dynamics (Stadler et al., 2013; Louca and Pennell, 2020a). Importantly, such inference relies on statistical identifiability of the time-dependent birth-death models, which is one of the conditions that guarantee that the estimated rates approach their true values as the tree size increases.

Kubo and Iwasa (1995) initially showed that time-dependent birth-death models are unidentifiable for reconstructed trees, that is, multiple pairs of birth and death rates have the same probability of producing a reconstructed tree. Despite the age of Kubo and Iwasa’s result, and the continuing popularity of these models, widespread concern (Pagel, 2020) about this unidentifiability issue was only recently prompted by the work of Louca and Pennell (2020b). The major implication of the unidentifiability is that we cannot estimate the two rate functions without placing additional constraints (Morlon et al., 2022) on one of them. Crucially, it has been shown that the number of congruent pairs of birth and death rates, — the rates that produce the same distributions of reconstructed trees, in the absence of such constraints is infinite (Kubo and Iwasa, 1995; Louca and Pennell, 2020b). This rules out manual parameter selection from the class of equally likely parameter pairs as a mitigating strategy.

The unidentifiability of the time-dependent birth-death models have been at least partially alleviated by Legried and Terhorst’s findings that piecewise constant, and subsequently, piecewise polynomial birth-death models, are identifiable for reconstructed trees (Legried and Terhorst, 2022, 2023). However, no such result for other classes of models such as those involving exponential growth or decay has been obtained.

In many applications, samples are obtained at different points in time, for example, when fossils are included in the analysis or when heterochronous sampling of pathogen genomic material is employed. Modelling such sampling through time is very important to avoid bias in the rate estimates (Stadler, 2010). For this reason, an additional parameter, sampling rate, was added to the birthdeath process, introducing a birth-death sampling process (Morlon et al., 2011; Stadler, 2010; Stadler et al., 2013; Gavryushkina et al., 2014; MacPherson et al., 2021). Early applications of this process to inference assumed that lineages are removed from the process immediately after sampling, which might be the case for some infectious diseases (e.g. HIV), where the diagnosis typically prevents a patient from transmitting the disease further. Louca et al. (2021) showed that this time-dependent birth-death sampling model is again unidentifiable when we consider reconstructed trees.

Removing lineages immediately after sampling is a strong and unrealistic assumption (Andréoletti and Morlon, 2023) for many other processes, e.g. fossilisation or transmission process for highly transmittable diseases where isolation is not performed on the diagnosis, such as influenza or coronavirus. A birth-death sampling model where sampling does not imply removing the lineage from the process is called the fossilised birth-death (FBD) model (Stadler, 2010; Heath et al., 2014). This model is widely used to estimate fossilisation rates and date species phylogenies (Stadler et al., 2018; Wright et al., 2022), however, the identifiability of the time-dependent rates for the FBD model has not been established (Andréoletti and Morlon, 2023).

In this paper, we show that the time-dependent FBD model is identifiable for arbitrary rate functions with strictly positive sampling rate. Although this does not imply that it is practically possible to estimate arbitrary rates from finite phylogenies, it provides a strong theoretical basis for using the FBD model to estimate speciation, extinction, and sampling parameters from phylogenies, and eliminates the requirement that the rates are restricted to be piecewise-polynomial. We demonstrate an important relationship between the distributions of complete and reconstructed trees under the time-dependent FBD model. We also show that the time-dependent birth-death sampling model (MacPherson et al., 2021), in which the removal probability is another unknown time-dependent parameter, is unidentifiable. That is, if the status of a sample after sampling is unknown and likely changes with time, we are again unable to estimate the birth and death rates.

Further, we use normalised deterministic lineages through time (nLTT) curves to demonstrate how the number of lineages in the reconstructed trees is affected by allowing sampled ancestors, that is, allowing species sampled in the past to be ancestors of younger species, which is the case when sampling does not imply removal. Finally, we discuss the implications of our results for data analysis.

## 2 Materials & Methods

### 2.1 Preliminaries

Here we consider a time-dependent fossilised birth-death (FBD) process (Stadler, 2010; MacPherson et al., 2021; Heath et al., 2014; Nee et al., 1994). The time in this process is usually set backwards, and the process runs from the time of origin *t*_0_ > 0 until the present (time zero). It starts with a single lineage, and all lineages arising in the process independently bifurcate, die, and/or are sampled through time with time-dependent Poisson rates *λ*(*t*), *µ*(*t*), and *ψ*(*t*). These rates are lineage-independent, i.e. all lineages at the same time point will have the same birth, death, and sampling rates. At the present time, extant lineages are additionally sampled uniformly at random with probability *ρ*_0_, called the *sampling fraction*. This time-dependent FBD process is a special case of the time-dependent birth-death sampling process (MacPherson et al., 2021) in which there is an extra parameter: the *removal probability*, which is a probability of a lineage being removed from the process immediately after sampling. In the FBD process this probability is set to zero, reflecting that sampling does not influence whether lineages remain in the process.

All lineages created by such a process together with the marks for the sampling events form a phylogenetic tree, which we call a *complete* tree 𝒯 (Fig. 1). To write down the probability density of complete trees under the FBD process we need to introduce the following notation, that we keep consistent with MacPherson et al. (MacPherson et al., 2021). We assume that *ρ*_0_ = 1 when considering complete trees. Let *N*_0_ denote the number of present day taxa, each of which is sampled at present. The tree has *n* death events at times (*y*_1_, *y*_2_, … *y*_*n*_), *N*_0_ + *n* − 1 branching events at times 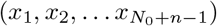, and *m* sampling events at times (*z*_1_, *z*_2_, … *z*_*m*_). The first branching event at *x*_1_ is called the root of the tree. A tree 𝒯 can be split into its discrete and continuous components and can be written as a pair 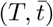 where *T* is the ranked ordered tree topology (Gavryushkin and Drummond, 2016; Steel, 2016; Gavryushkin et al., 2018), and 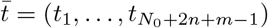 is the vector of times of branching, death, and sampling-through-time events with 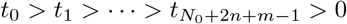.

**Figure 1:**
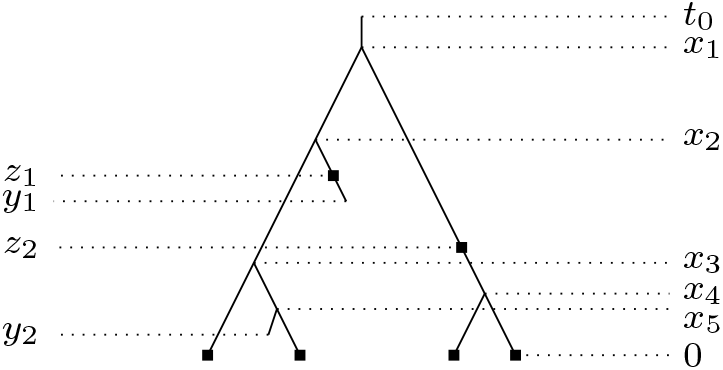
An example of a complete tree generated by a time-dependent Fossilised Birth-Death (FBD) process. The stem of the tree is denoted *t*_0_, and the present occurs at time 0. Birth events occur at times *x*_*i*_, death events at times *y*_*j*_, and samples at times *z*_*k*_.

In what follows, we assume that *λ*(*t*) ≥ 0, *µ*(*t*) ≥ 0, and *ψ*(*t*) ≥ 0 for all *t* ∈ [0, *t*_0_] and are Lebesgue-integrable on [0, *t*_0_]. Following the well described derivation (see, e.g. Stadler, 2010; Morlon et al., 2011; MacPherson et al., 2021), we obtain the probability density function for the complete trees under the time-dependent FBD process:

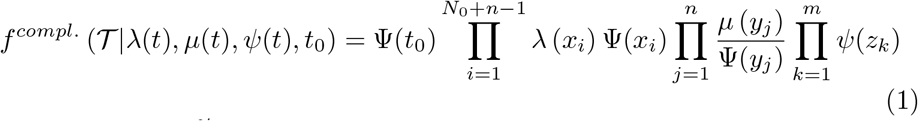

where 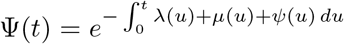. Each node in the tree contributes to the probability density function as the rate of the event (birth, death, or sampling) at the corresponding time, and each edge contributes the exponential term, Ψ (*t*), reflecting that no events took place along them. SI Figure S1 illustrates the parts of a tree that correspond to the exponential of the integral terms.

Note that it is common in the literature to use the term “likelihood” for the probability density function even if it is not considered as a function of parameters. In this paper, we prefer to use the term “probability density function”.

In a complete tree, all speciation, extinction, and sampling events (through time and at present) are observed. However, in practice we are not able to observe (or infer) the complete tree, and therefore parameter estimation relies on the *reconstructed* tree. The reconstructed tree is a tree induced by the sampled nodes and is shown in Figure 2b given the complete tree in Figure 2a.

**Figure 2:**
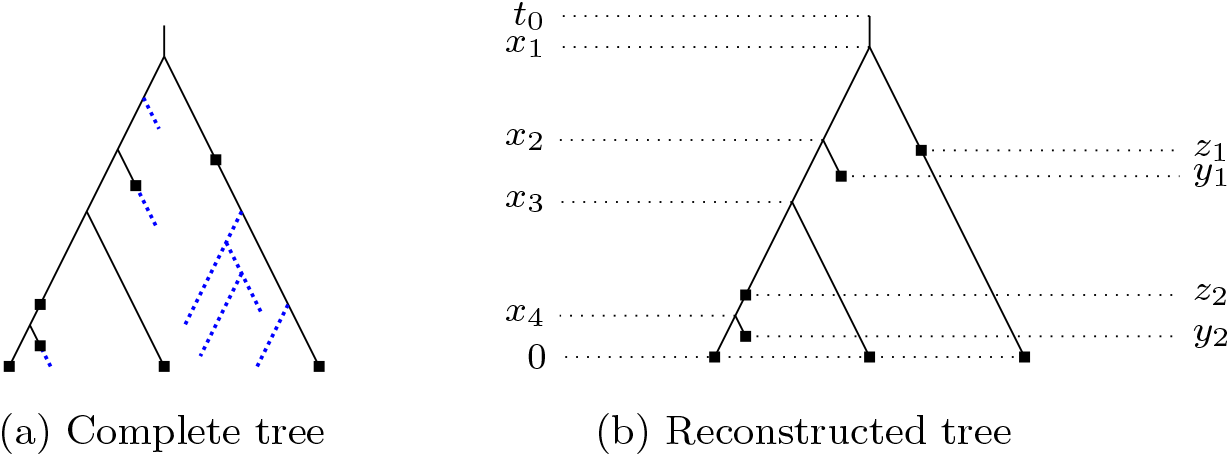
An example of a complete tree (a) and the corresponding reconstructed tree (b) obtained from the complete tree by trimming unobserved lineages (blue, dotted). The unobserved lineages are the lineages that do not terminate with samples (solid squares). In the resulting reconstructed tree, there are three types of nodes: branching nodes at times (*x*_*i*_), tip samples (*y*_*i*_ and present time), and sampled ancestors (*z*_*i*_).

We now introduce a notation for reconstructed trees which is similar to that used in the complete tree case. The sampling at present probability *ρ*_0_ can now take any value between zero and one. The reconstructed tree has *M*_0_ sampled extant species, *M*_0_ + *n* − 1 branching events, *n* sampling events that result in termination of lineages in the tree (tips), and *m* remaining sampling events that lie within lineages of reconstructed tree and are called *sampled ancestors*. We denote the times of the branching events as 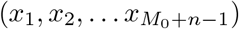, the times of the terminal sampling events as (*y*_1_, *y*_2_, … *y*_*n*_), and the times of the remaining sampling events as (*z*_1_, *z*_2_, … *z*_*m*_) (Fig. 2b).

To derive the probability density function for reconstructed trees, we need to define function *E*(*t*), which is the probability that a lineage alive at time *t* does not leave sampled descendants. It was shown (MacPherson et al., 2021) that this function is the solution to the following differential equation:

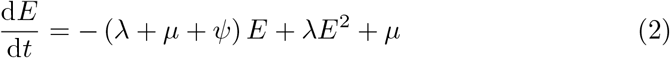

with initial condition *E*(0) = 1 − *ρ*_0_. The following proposition establishes the conditions on the rate functions which guarantee that the solution to this equation exists and is unique.

**Proposition 2.1**. Equation (2) has a unique extended solution on [0, *t*_0_] if the following conditions are satisfied:

- *λ* ≥ 0, *µ* ≥ 0, and *ψ* ≥ 0 are Lebesgue-integrable on [0, *t*_0_],
- 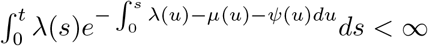 for *t* ∈ [0, *t*_0_], and
- either *ρ*_0_ > 0 or *ψ*(0) > 0 and *λ, µ*, and *ψ* are continuous at zero.

*Proof*. See SI Section S2

In what follows, whenever reconstructed trees are considered, we assume that *λ, µ*, and *ψ* satisfy these conditions.

The probability density function for reconstructed trees (MacPherson et al., 2021) is:

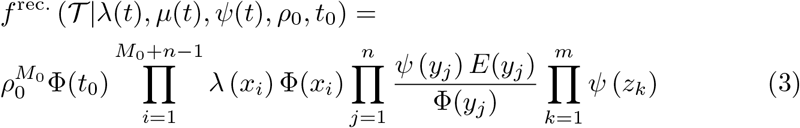

where 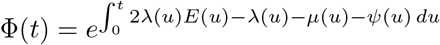. Note that 1−*E* (*t*_0_) is the probability that at least one lineage was sampled either in the past or the present (Morlon et al., 2011). Then the probability density function for reconstructed trees, conditioning on the event of sampling at least one lineage (*S*), is:

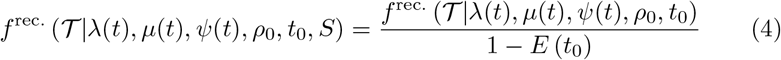

Consider a parametric statistical model, which is a class of distributions *P*_*θ*_ across a set of parameters Θ: 𝒫_Θ_ = {*P*_*θ*_ : *θ* ∈ Θ}.

**Definition 2.1** (Identifiability). A statistical model 𝒫_Θ_ = {*P*_*θ*_ : *θ* ∈ Θ} is identifiable if *θ* → *P*_*θ*_ is injective; that is, for all *θ*_1_, *θ*_2_ ∈ Θ, we have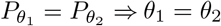. Two probability density functions 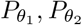 are considered equal if the set of points where they differ has Lebesgue measure zero.

The parameters *θ* of the FBD model are (*λ, µ, ψ, ρ*_0_, *t*_0_). We consider two models for which we show identifiability. The complete tree model is described by probability densities 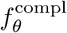. defined by equation (1) with restrictions on the parameters formulated in Theorem 3.1 that define set Θ_compl._. The reconstructed tree model is described by probability densities 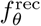. defined by equation (3) and has a slightly different set of allowable (with restrictions from theorem 3.2) parameters Θ_rec._.

### nLTT code implementation

In unidentifiable models (birth-death or birth-death sampling with removal at sampling cases) there are sets of birth, death, and sampling rates for which the probability density functions and therefore reconstructed tree distributions are identical (Louca and Pennell, 2020b; Louca et al., 2021). Such parameters form congruence classes, where any pair of parameters are congruent if they produce the same distributions of reconstructed trees.

To demonstrate congruent scenarios Louca and Pennell (2020b) and Louca et al. (2021) used deterministic lineage through time (dLTT) curves. This curve is the deterministic limit of the number of observed lineages at times *t* generated in the stochastic birth-death sampling processes (Kubo and Iwasa, 1995). The stochastic process describes the number of lineages present, given diversification and sampling events which are realisations of a birth-death sampling process. Deterministic calculations assume that the timing of events is solely determined by the birth, death, and sampling rates with no random component. dLTT curve also corresponds to the expected number of lineages in the reconstructed trees generated by the stochastic birth-death sampling process (SI S4.2). Therefore, pairs of rates that produce different dLTT curves also produce different distributions of reconstructed trees.

Here we follow Louca et al. (2021) who considered a normalised lineage through time (nLTT) curve, which is a dLTT curve normalised by its area under the curve (Janzen et al., 2015). Obviously, different nLTT curves imply different dLTT curves. To plot nLTT curves Louca et al. (2021) used the R package *castor* (Louca and Doebeli, 2018). They randomly selected birth, death, and sampling rates, which were generated under either an exponential or an Ornstein-Uhlenbeck stochastic process. Here we either randomly select exponential rates as in Louca et al. (2021) or manually select linear rates to avoid technical difficulties with obtaining congruent scenarios with positive rates. Given the selected rates *θ*_1_ = (*λ*_1_, *µ*_1_, *ψ*_1_), we use the castor function **congruent hbds model** and calculations based on the pulled diversification rate to obtain congruent rates *θ*_2_ = (*λ*_2_, *µ*_2_, *ψ*_2_). Then we calculate deterministic values using the **simulate deterministic hbds** function by generating a tree with between 100,000 and 200,000 tips, given the rates (*θ*_1_ or *θ*_2_) and sampling fraction *ρ*_0_ = 0 to achieve a good approximation of the nLTTs. Rather than utilising the removal probability *r*, the package functions use the retention probability 𝒦 = 1 − *r*, which we specify accordingly.

We use a fork of *castor v*.*1*.*7*.*11* suitable for calculating nLTT curves for trees with sampled ancestors. This is available at https://github.com/bioDS/ castor. The R code from Louca et al. (Louca et al., 2021) was used as a basis for the simulations. The deterministic parameter calculation code for FBD trees is available at https://github.com/bioDS/FBD-dLTTs.

## 3 Results

### 3.1 Identifiability of the time-dependent FBD model

Here we show the identifiability of the time-dependent FBD model parameters from complete trees.

#### Theorem 3.1

Suppose *t*_0_ is fixed and *λ*(*t*) ≥ 0, *µ*(*t*) ≥ 0, and *ψ*(*t*) ≥ 0 are Lebesgue-integrable on [0, *t*_0_]. Then the time-dependent FBD complete tree model (the probability density function for this model is defined by equation 1) is identifiable.

*Proof*. The key ingredient of the proof of the theorem is the following lemma:

#### Lemma 3.1

Assume 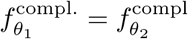. almost everywhere, then the following equalities hold:

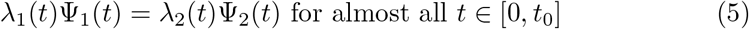

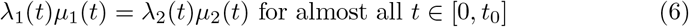

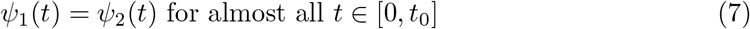

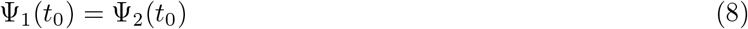

*Proof*. Equalities (5)–(7) are obtained by considering trees that differ by one event as follows. To show (5), we note that the densities should be equal for almost all trees with *s* edges and *s*_*e*_ extant edges, and also for almost all trees with *s* + 1 edges and *s*_*e*_ + 1 extant edges. Given a pair of trees in which one tree is obtained from the other by attaching an additional edge at time *t* (Figs. 3a and 3b), their densities differ by one term, namely, *λ*(*t*)Ψ(*t*), which implies equality (5).

**Figure 3:**
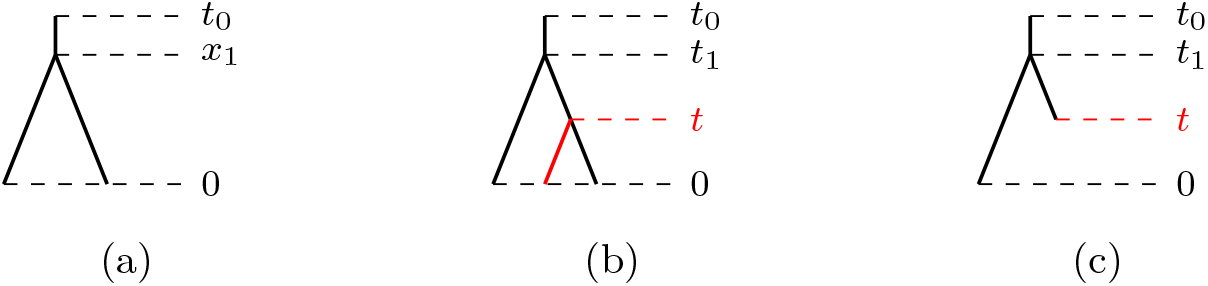
Tree modifications used in the proof of Lemma 3.1. Subfigure (a) depicts the original tree, while Subfigure (b) shows the addition of an extant edge born at time *t* and Subfigure (c) shows the contraction of an extant edge such that it dies at time *t*.

By considering pairs of trees in which one edge switches from being extant to going extinct at time *t* (Figs. 3a and 3c), we obtain 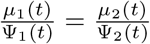,which together with (5) implies (6). Similarly, considering trees with an additional sampling event, we obtain (7). Since all the individual terms except for Ψ(*t*_0_) are equal in the probability density functions of all trees under *θ*_1_ and *θ*_2_, we conclude that the Ψ(*t*_0_) terms should also be equal (8). The detailed proof can be found in SI section S3.2.

We now finish the proof of the theorem as follows. Denote 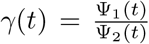. Equalities (5)-(7) imply that *γ* must satisfy the following differential equation:

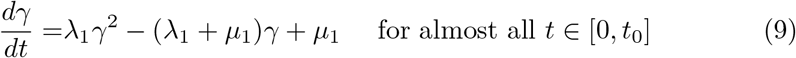

Definition of *γ* and equality (8) implies the boundary condition *γ*(0) = *γ*(*t*_0_) = 1. *γ*(*t*) = 1 is a solution to this equation, which must be unique (SI section S3.3). Therefore, *λ*_1_(*t*) = *λ*_2_(*t*) and *µ*_1_(*t*) = *µ*_2_(*t*) for almost all *t* ∈ [0, *t*_0_].

#### Theorem 3.2

Suppose *t*_0_ is fixed, parameters *λ, µ, ψ* and *ρ*_0_ satisfy the conditions of Proposition 2.1, and *ψ*(*t*) > 0 for almost all *t* ∈ [0, *t*_0_]. Then the time-dependent FBD reconstructed tree model (the probability density function is defined by equation 3) is identifiable.

*Proof*. We prove this theorem by showing that there exists a one-to-one transformation of the FBD parameters (Eq. (10), Lemma 3.2) such that the distribution of the reconstructed trees produced by the process with original parameters and conditioned on sampling of at least one lineage is the same as the distribution of complete trees produced by the FBD process with transformed parameters (Lemma 3.3).Then we apply Theorem 3.1 to complete the proof.

Consider the following parameter transformation of original rates to *pulled* rates:

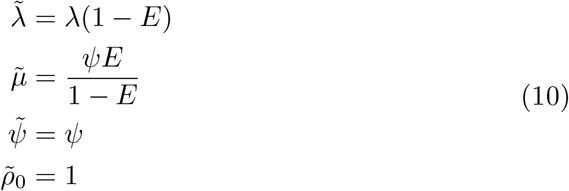

where *E*(*t*) is defined in Equation (2). Note that since *ψ*(*t*) > 0 we have *E*(*t*) < 1 for all *t* ∈ (0, *t*_0_] (SI Section S2). Then if *ρ*_0_ = 0, implying *E*(0) = 1, we set 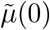 to zero. Thus, all the rates are well-defined for all *t*.

We say that two sets of rates *λ*_1_(*t*), *µ*_1_(*t*), *ψ*_1_(*t*) and *λ*_2_(*t*), *µ*_2_(*t*), *ψ*_2_(*t*) are different if at least one of the rates (*λ, µ*, or *ψ*) differ on at least a non-zero measure subset of [0, *t*_0_].

#### Lemma 3.2

Parameter transformation (10) is a one-to-one correspondence.

*Proof*. See SI Section S4.3.

#### Lemma 3.3

The distribution of reconstructed trees for the time-dependent FBD process, conditioned on sampling of at least one lineage (through-time or at present), is the same as the distribution of the complete trees under the time-dependent FBD process with pulled rates and a sampling probability of one. That is,

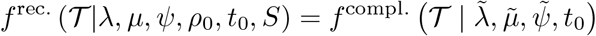

*Proof*. The proof follows from substituting the pulled rates into the probability density function for the complete trees (SI Section S4).

To finish the proof of the theorem suppose 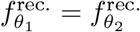.. Then 1 − *E*_1_(*t*_0_) = 1 − *E*_2_(*t*_0_), because 1 − *E*(*t*_0_) is the cumulative probability of all surviving trees. So 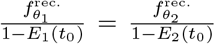.. Lemma 3.3 implies that 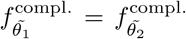.. Then it follows from Theorem 3.1 that 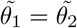, and from Lemma 3.2 that *θ*_1_ = *θ*_2_.

Note, that if *ψ* is equal to zero on some time interval within [0, *t*_0_] then only birth or death events can happen at this time interval. Therefore within this time interval, the process can be described by only one pulled birth rate as shown in (Louca and Pennell, 2020b) and the model becomes unidentifiable. In other words, by choosing identical birth, death and sampling rates whenever *ψ* is greater than zero and congruent birth and death rates (as defined for unidentifiable birth-death models) whenever *ψ* is zero we obtain congruent rates for the reconstructed FBD model.

### 3.2 Unidentifiability of the birth-death model with removal at sampling

In this section we address the identifiability problem for a time-dependent birth-death sampling model with removal at sampling (MacPherson et al., 2021). This model is defined by a time-dependent FBD process where sampled lineages are removed from the process after sampling with time-dependent removal probability *r*(*t*), where 0 ≤ *r*(*t*) ≤ 1.

For this model, we can also define pulled rates:

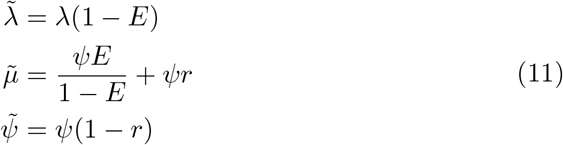

where *r* ≠ 0 on some non-zero measure subset of [0, *t*_0_]. Similarly to the FBD case, the probability density function of the reconstructed trees with original rates is the same as the probability density function of complete trees under the model with pulled rates (see SI section S5.2).

Now, for any given *λ* > 0, *µ* ≥ 0, *ψ* > 0, 1 ≥ *r* ≥ 0, *ρ*_0_ ∈ (0, 1], and for any alternative rate *ψ*^*^ > 0 (here we also assume *λ, ψ*, and *ψ*\* are differentiable almost everywhere on [0, *t*_0_]) we define

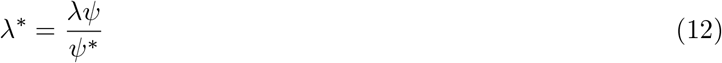

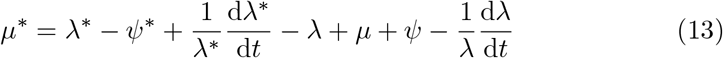

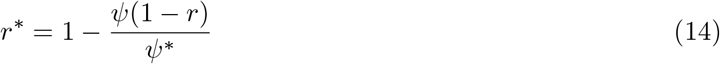

Equation (14) implies 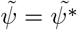. Louca et al. (Louca et al., 2021) showed that (12) and (13) imply:

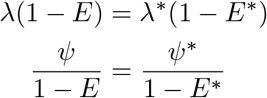

Therefore we have 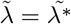.Noting that 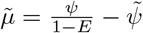 we have 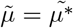.

Thus for a given model with parameters *θ* and an arbitrary *ψ*^*^ distinct from *ψ* we obtained another model defined by distinct parameters *θ*^*^ for which the pulled rates are equal and therefore the probability distributions over reconstructed trees are also equal. Given that we have found distinct parameter sets that produce the same probability distributions, the following theorem holds:

#### Theorem 3.3

Suppose *t*_0_ is fixed, parameters *λ, µ, ψ* and *ρ*_0_ satisfy the conditions of Proposition 2.1, *ψ*(*t*) > 0 for almost all *t* ∈ [0, *t*_0_], *λ* and *ψ* are differentiable almost everywhere on [0, *t*_0_], 0 ≤ *r*(*t*) ≤ 1, and *r* ≠ 0 on an non-zero measure subset of [0, *t*_0_]. Then the time-dependent birth-death sampling model with removal at sampling is unidentifiable for reconstructed trees.

In the above theorem, we assumed a restricted set of parameters for which the unidentifiability holds. For any wider parameter set that contains this restricted set, the model remains unidentifiable. However, if stricter or non-overlapping conditions apply, the model may become identifiable, e.g. fixing *r* to zero (FBD model identifiability) or fixing *ψ* to zero and considering piecewise constant or piecewise polynomial birth and death rates (Legried and Terhorst, 2022, 2023). Furthermore, if we fix the removal probability to a function that is not equal to one then given that the pulled rate transformation 11 is one-to-one (SI section S5), we obtain the following corollary:

#### Corollary 3.1

Suppose *t*_0_ is fixed, parameters *λ, µ, ψ* and *ρ*_0_ satisfy the conditions of Proposition 2.1, *ψ*(*t*) > 0 for almost all *t* ∈ [0, *t*_0_], 0 ≤ *r*(*t*) ≤ 1, and *r*(*t*) is a fixed function such that *r*(*t*) ≠ 1 almost everywhere on [0, *t*_0_]. Then the time-dependent birth-death sampling model with fixed removal at sampling probability is identifiable for reconstructed trees.

### 3.3 nLTT curves

Louca et al. (2021) illustrated the congruency of the birth-death model by comparing the nLTT curves produced using different congruent rate functions. We follow this methodology to assess deterministic properties given particular speciation, extinction, and sampling rates, in our case under the FBD process. We use two sets of rate functions, *θ*_1_ and *θ*_2_. Given *θ*_1_ = (*λ*_1_, *µ*_1_, *ψ*_1_), *θ*_2_ = (*λ*_2_, *µ*_2_, *ψ*_2_) is chosen such that *θ*_1_ and *θ*_2_ result in the same nLTT curve under the time-dependent birth-death sampling model with a removal probability *r* = 1. We then use *θ*_1_ and *θ*_2_ to produce nLTT curves for scenarios under the process with smaller removal probabilities, specifically *r* = 0.5 and *r* = 0, which allow for sampled ancestors.

Figure 4 illustrates the diversification rates and deterministic curves for a scenario with linear diversification rates for *θ*_1_, while Figure 5 depicts a scenario with exponential forms for *θ*_1_. Note that restricting *θ*_1_ to linear or exponential rates does not place the same restriction on *θ*_2_. We observe that the introduction of sampled ancestors for the same speciation, extinction, and sampling rates affects the nLTT curve, particularly near its peak. As time passes and sampling events occur without removing lineages, we expect the total number of speciation, extinction, and sampling events to increase with the number of lineages, given the same lineage-independent rates.

**Figure 4:**
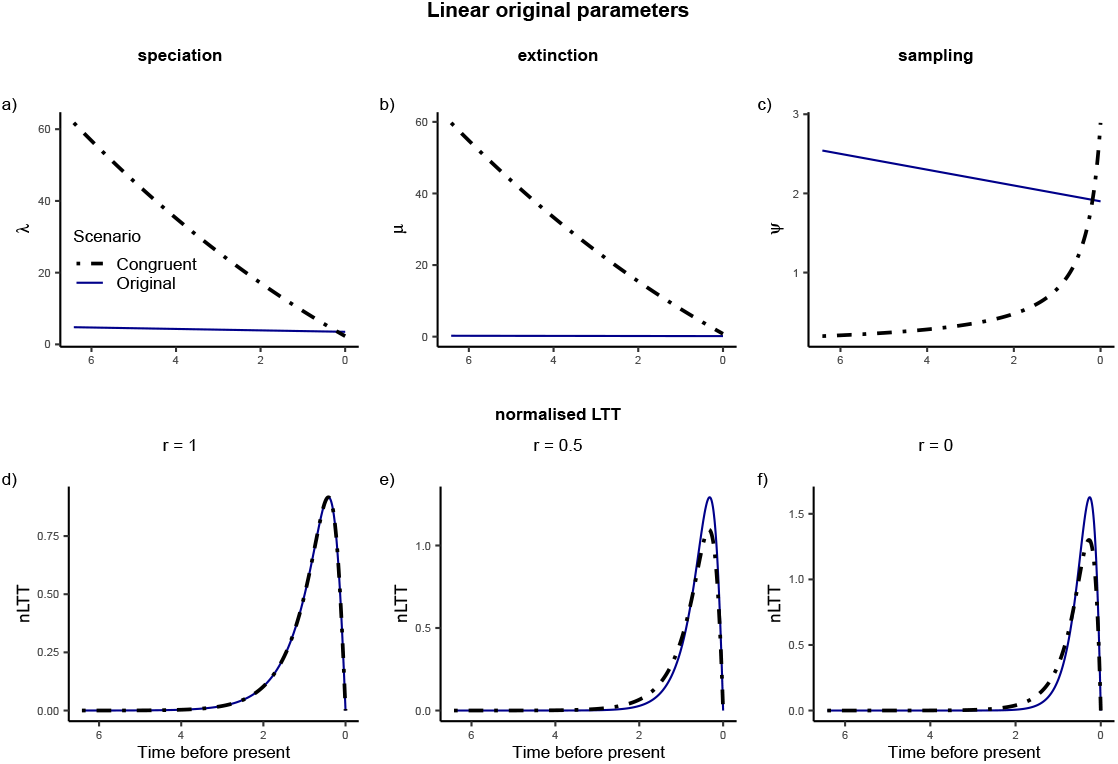
A set of congruent scenarios where *θ*_1_ has linear rates. Subfigures a–c) show the parameters of *θ*_1_ and *θ*_2_ where *θ*_1_ = (0.2*t* + 3.5, 0.015*t* + 0.15, 0.1*t* + 1.9). Subfigures d–f) illustrate that when sampled ancestors are permitted, *θ*_1_ and *θ*_2_ no longer produce identical nLTT curves.

**Figure 5:**
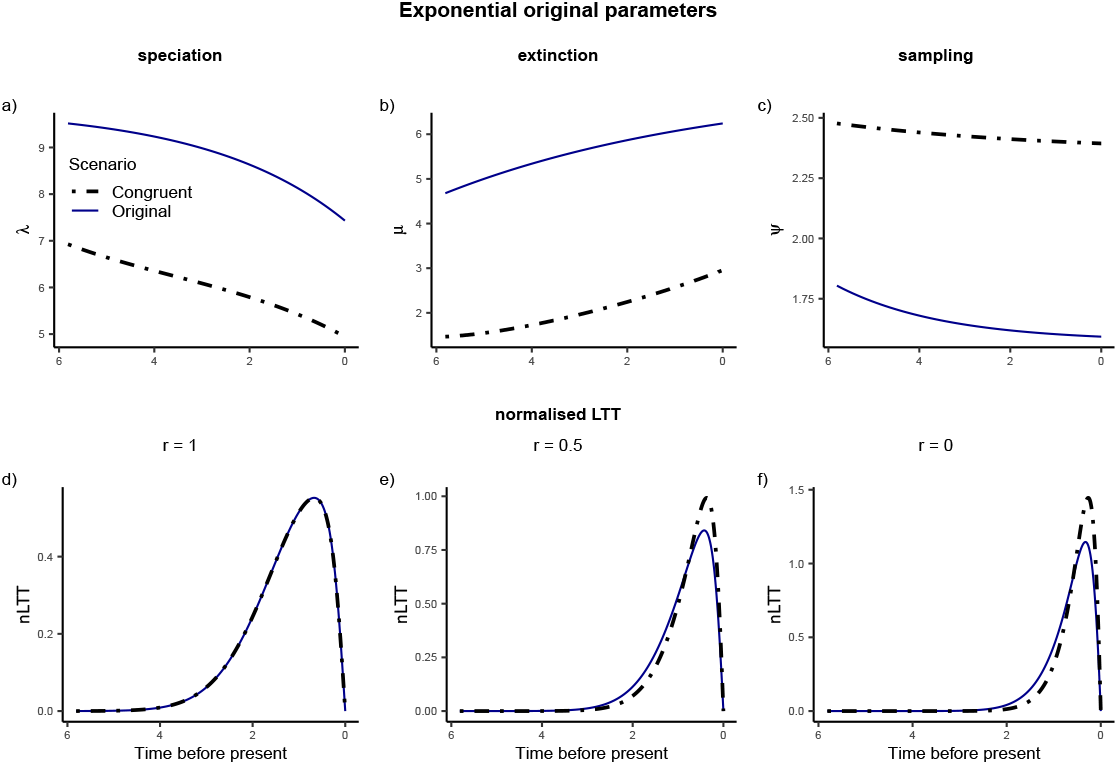
A set of congruent scenarios where *θ*_1_ has exponential rates which were randomly selected. Subfigures a–c) show the parameters of *θ*_1_ and *θ*_2_ where *θ*_1_ ≈ (9.84 − 2.41*e*^−0.35*t*^, 7.16 − 0.92*e*^0.17*t*^, 1.57 + 0.02*e*^0.43*t*^). Subfigures d–f) illustrate that when sampled ancestors are permitted, *θ*_1_ and *θ*_2_ no longer produce identical nLTT curves.

While using a removal probability of less than one results in different nLTT curves for scenarios that were previously congruent under *r* = 1, the difference in the nLTT may be small for all times considered. An example of this is shown in Figure 6. This small difference between nLTT curves is observed in scenarios where the rate of sampling is low compared to the total tree length. In such cases, there are relatively few time points at which a lineage may be removed due to sampling, and thus the number of lineages in trees is largely unaffected by the removal probability.

**Figure 6:**
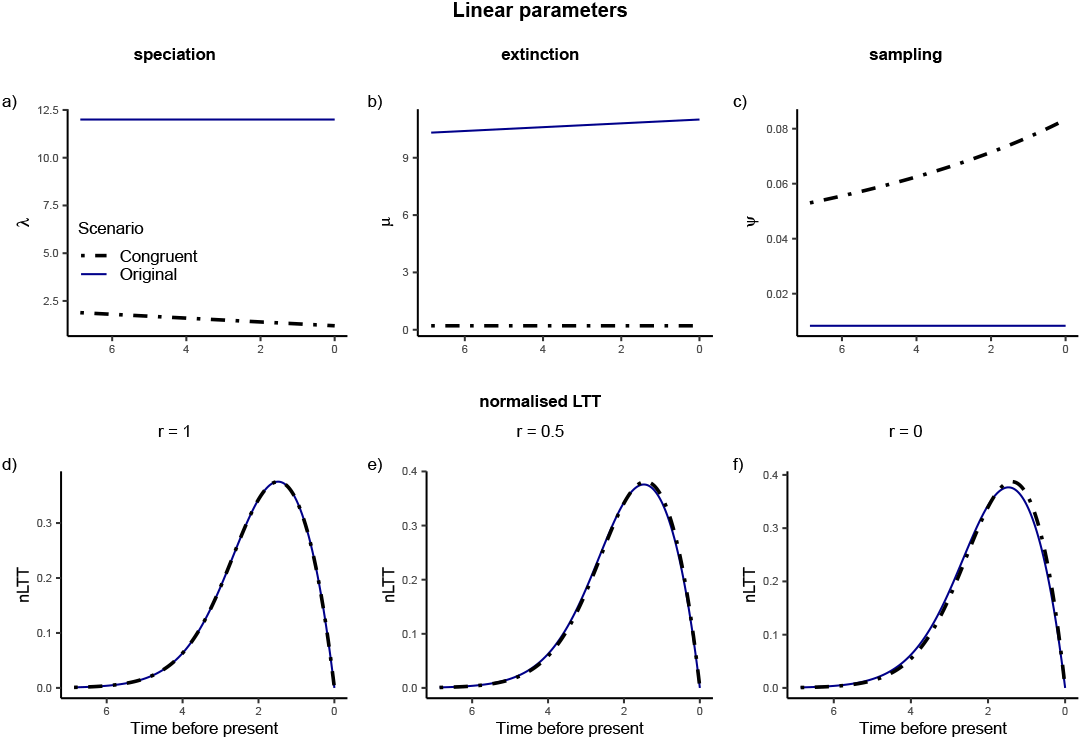
A set of congruent scenarios, where *θ*_1_ has linear rates. Subfigures a–c) show the parameters of 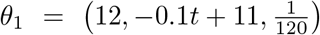 and 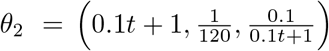 Subfigures d–f) illustrate that although reducing the removal probability results in different nLTT curves for *θ*_1_ and *θ*_2_, the difference in density is small for all times considered.

A direct comparison of nLTT curves under different linear removal probabilities is shown in Figure 7. In this example scenario, the speciation, extinction and sampling rates are fixed. The linear removal probability functions with different slops reflect scenarios ranging from rapid growth of removal probability with time to constant removal probability. We observe that the steeper the removal probability function, the greater the deviation in the nLTT curve compared to the case where the removal probability is constant (shown as a solid line).

**Figure 7:**
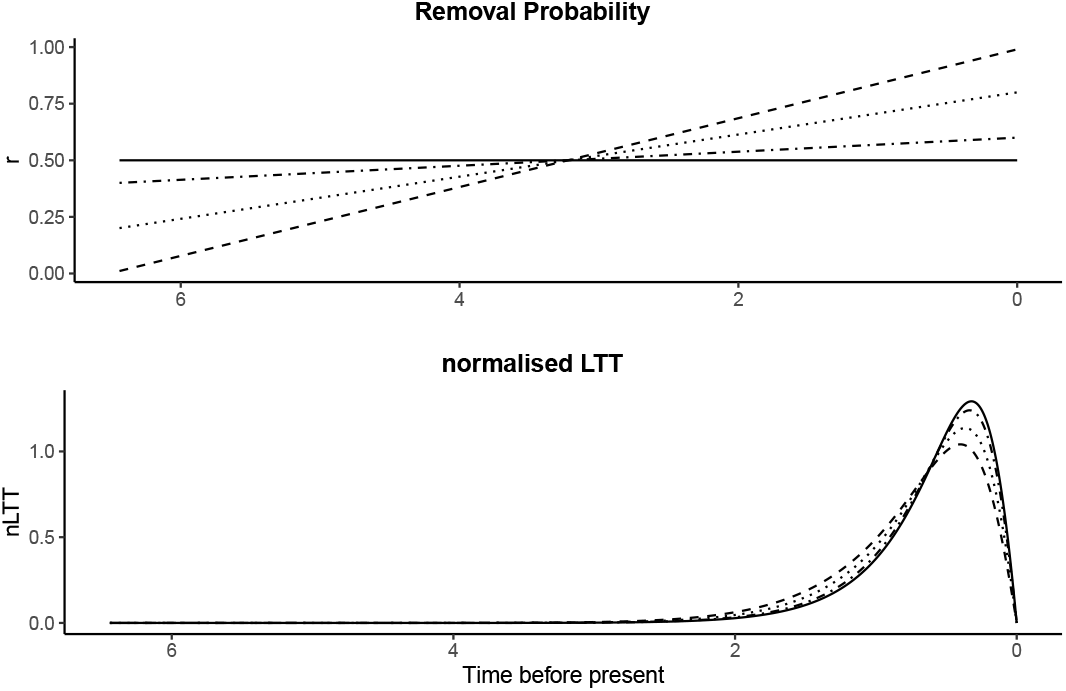
An example of differences in nLTT curves caused by changing the removal probability *r* while keeping speciation, extinction and sampling rates fixed: *θ* = (0.2*t* + 3.5, 0.015*t* + 0.15, 0.1*t* + 1.9).

## 4 Discussion

In this study, we have shown that the time-dependent FBD model is identifiable and therefore eliminated the concern related to the use of such models for parameter inference (Pagel, 2020) caused by the previous unidentifiability results. Given the identifiability, the FBD model is an absolutely suitable model for inferring speciation and extinction rates from dated phylogenies, as well as epidemiological parameters in scenarios where removal upon diagnosis is not implied.

Previous studies showed identifiability of the birth-death model without sampling for piecewise constant and piecewise polynomial rates. The FBD model is identifiable in an arbitrary class of rate functions. While a piecewise constant model can be defined to approximate an unidentifiable model arbitrarily closely (Legried and Terhorst, 2022), it is unclear how much a particular piecewise constant model that maximises the likelihood function deviates from the true non-piecewise constant set of rates. Although computational methods do not allow us to infer arbitrary functions, and we still need to restrict the form of the estimated rates, we can now directly model, for example, exponential growth, which would reduce the number of estimated parameters and required tree size compared to piecewise constant or polynomial approximations.

Previous implementations of the FBD model have relied on simulation studies to show identifiability (Gavryushkina et al., 2014); our theoretical results make this time-consuming step of investigating parameter identifiability via simulation unnecessary, whether one wishes to use constant rates, piecewise-constant, piecewise-polynomial, or more complex scenarios. Such simulations can still be useful for assessing the precision of estimated parameters, robustness of the model to assumptions violation or validation of software implementations.

It is important to note, that although the FBD model is theoretically identifiable, a given dataset may not be of a sufficient size for rate selection (Morlon et al., 2022; Legried and Terhorst, 2023). While theoretically unidentifiable models cannot be resolved with any amount of data (Louca and Pennell, 2020b), there is no fixed finite quantity of data for theoretically identifiable scenarios which guarantees a desired precision of the inferred rates. Small trees or samples may lead to wide confidence or credible intervals for parameters such as extinction rate even in the case when the true rates have simple forms (Morlon et al., 2022). A further caveat to applying theoretical identifiability results is that they unrealistically rely on knowing the true phylogeny without an error. With limited molecular, morphological, or other data, simplified evolutionary models and otherwise non-perfect inference methods, knowing the true phylogeny without an error is not possible.

The results of our study show the importance of including fossils when estimating speciation and extinction rates. The general case when only a dated phylogeny is available is consistent with many congruent rates. Once fossils are placed on the phylogeny, the congruence class collapses to a single set of rates that are consistent with the given sampled ancestor phylogenetic tree (a tree in which fossil samples can be sampled ancestors, Gavryushkina et al., 2014). Including sampled ancestor fossils has been shown to be very important in reducing biases in estimated extinction rates (Gavryushkina et al., 2014; O’Reilly and Donoghue, 2019). Here, we have shown that allowing fossils to be sampled ancestors is crucial for theoretical identifiability of the diversification rates.

It is intuitive that allowing removal probabilities of less than one provides more information about the sampling rate, which in turn helps determine the birth and death rates. As a lineage under the time-dependent FBD model may survive being sampled, there is the potential for each lineage to be sampled on multiple occasions. These additional sampling events define the time intervals between sampling on continuing lineages, which are informative for determining the sampling rate. Whereas if sampling guarantees removal, the sampling rate is confounded by the unknown birth and death rates. It becomes impossible to distinguish between scenarios with low sampling rate and many unobserved lineages and high sampling rate with few unobserved lineages.

The ability to infer the sampling rate itself, even if the sampling scheme was not constant through time, is an additional benefit of the FBD model. Inferred fossilisation rates could for example be further used in consecutive studies as prior information.

As we demonstrated in the nLTT examples, the sampling rate has to be high enough to introduce noticeable differences in the distributions of the reconstructed trees and therefore enable precise maximum likelihood or Bayesian inference of the rates. Our identifiability finding cannot be applied to scenarios which allow past time periods with no probability of sampling lineages. The proven one-to-one correspondence between complete and reconstructed trees relies upon a non-zero past sampling rate to determine original rates from pulled rates. Given that species’ physical structure or habitat may make them unsuitable for fossilisation (Shaw et al., 2021), the FBD model might not be suitable for scenarios requiring *ψ* (*t*) = 0 for significantly long periods of time. However if the birth and death rates are piecewise polynomial where *ψ* is equal to zero we can apply the results of Legried and Terhorst (Legried and Terhorst, 2022, 2023) to obtain the identifiability of the reconstructed FBD model.

It is important to note, that although the inclusion of fossils helps identify the speciation and extinction rates, the age uncertainty and biased sampling associated with fossil data could still propagate to the speciation and extinction rates estimates. Modelling fossil age uncertainty and using stratigraphic ranges of species as opposed to single fossil representatives (Stadler et al., 2018) could improve the accuracy of credible intervals and reduce the bias.

The identifiability of the time-dependent FBD model arises from allowing sampled fossils to remain in the process (removal probability *r* fixed to zero) and no longer holds once the removal probability is set to one (Louca et al., 2021), as has been previously shown for piecewise constant rates (Gavryushkina et al., 2014). Here we have also shown that the time-dependent birth-death sampling model remains identifiable, provided *r* is fixed to a function not equal to one. Such a model could be used in epidemiological studies for diseases where diagnosis does not imply that the patient prevents new transmissions, for example, influenza or coronavirus (where isolation strategies have not been applied). However, if one wants to estimate the removal probability from the data, the model becomes unidentifiable. In this case, one can impose strong prior assumptions on the rates (Morlon et al., 2022) or explore congruence classes for patterns that are conserved within them (Höhna et al., 2022; Kopperud et al., 2023).

While we have considered the impact of different fixed constant and linear removal probabilities on nLTT curves in several examples, the question of how misspecified removal probabilities impact the inference is yet to be explored. The difference in nLTT curves for different linear removal probabilities (Figure 7) demonstrates how this parameter affects the distribution of reconstructed trees, highlighting that a misspecified fixed removal probability could potentially influence diversification rate estimates. Potential avenues of inquiry include describing how the estimates of selected parameter rates are affected by the magnitude of error in removal probabilities.

By extending pulled rates introduced for birth-death sampling models (Louca et al., 2021) to the case of the FBD model, we also showed that the distributions of reconstructed and complete trees generated by related FBD processes are equivalent. This equivalence provides a different view on the nature of reconstructed trees that can be seen as complete trees generated by an FBD model with adjusted or pulled rates, as was previously noted by Louca et al. (2021) for trees without sampled ancestors. The original birth rate is adjusted by the probability of the newly born lineage to be observed. The original sampling rate is split in two: pulled death rate and pulled sampling rate, depending on the type of the occurred sampling event. Events that are followed by lineage disappearance from the observer become death events and events that are followed by further sampling events (sampled ancestors) become sampling events.

Despite the identifiability of the FBD model in the arbitrary class of rates, the inference methods require fixing the class of inferred rates. An avenue for future research would be to quantify how well different classes of functions that are available for inference (e.g., the widely used piece-wise constant functions) approximate the true rates of different function classes which imply identifiability of the corresponding FBD model.

Another direction for future research is to investigate identifiability for variations of birth-death models that account for lineage specific birth and death rates (Maddison et al., 2007), age-dependent rates (Hagen et al., 2015), or different speciation types (Stadler et al., 2018). While the identifiability of the pure birth multi-type model with some restrictions on the transition rates has been recently shown (Dragomir et al., 2023), the identifiability of the parameters in other models has not yet been generally established.

Finally, we would like to emphasise that this study completely answers the question of the time-dependent reconstructed FBD model identifiability. We have shown the identifiability for arbitrary rate functions (subject to the conditions that guarantee the probability distribution over the reconstructed trees can be defined) as long as the sampling rate is strictly positive. The last requirement is essential for identifiability, because allowing *ψ* to be equal to zero produces congruent scenarios and therefore implies unidentifiability.

In conclusion, the FBD model is an excellent candidate for modelling evolutionary processes. It shares computational conveniences of other birth-death models, describes sampling process with more realistic assumptions, and as we have shown here, possesses an essential statistical property of identifiability.

## Supplementary material

Drayd link: https://doi.org/10.5061/dryad.sj3tx96ch

## Acknowledgements

We thank Dmitry Berdinsky, Lars Berling, and Kerry Manson for their helpful discussions throughout this research.

## Author Contributions

ASG initiated the idea of proving identifiability. KT, TGV, AG, ASG extended the idea of transformed parameters to FBD case. KT, AG, ASG worked on the proofs. KT investigated nLTT plots. KT, AG, ASG wrote and edited the manuscript. TGV edited the manuscript.

