## Supplementary material for "The Fossilised Birth-Death Model is Identifiable"

### The Fossilised Birth-Death Model is Identifiable — Supporting Information

August 28, 2024

#### S1 Preliminaries

For convenience, we restate the notation used in the main article.

##### S1.1 Rate and present sampling probability parameters

The time dependent speciation, extinction, and sampling rates of the FBD process are denoted $\lambda(t)$, $\mu(t)$, and $\psi(t)$ respectively. The probability that an extant lineage is sampled, known as the sampling fraction, is $\rho_0$. We denote $\theta = (\lambda(t), \mu(t), \psi(t), \rho_0)$. The measure of a set $X$ is denoted $\boldsymbol{\mu}(X)$ to differentiate measures from extinction rates.

##### S1.2 Complete tree

The realisation of the time-dependent FBD process is a *complete* tree $\mathcal{T}$. This tree has $N_0$ present day taxa which are each sampled at present. The $n$ death events occur at times $(y_1, y_2, \ldots y_n)$, the $N_0 + n - 1$ branching events at $(x_1, x_2, \ldots x_{N_0+n-1})$, and the $m$ sampling events at $(z_1, z_2, \ldots z_m)$. The first branching event at $x_1$ is called the root of the tree. A tree $\mathcal{T}$ can be split in its discrete and continuous components and can be written as a pair $(T, \bar{t})$ where $T$ is the ranked ordered tree topology and $\bar{t} = (t_1, \ldots, t_{N_0+2n+m-1})$ is the vector of times of branching, death, and sampling-through-time events with $t_0 > t_1 > \cdots > t_{N_0+2n+m-1} > 0$.

The probability density function for the complete trees under the time-dependent FBD process is:

$$f^{\text{compl.}}(\mathcal{T}|\lambda(t), \mu(t), \psi(t), t_0) = \Psi(t_0) \prod_{i=1}^{N_0+n-1} \lambda(x_i)\,\Psi(x_i) \prod_{j=1}^{n} \frac{\mu(y_j)}{\Psi(y_j)} \prod_{k=1}^{m} \psi(z_k) \tag{1}$$

where $\Psi(t) = e^{-\int_0^t \lambda(u)+\mu(u)+\psi(u)\,du}$ (Figure S1). The function can be derived using the unified model approach from MacPherson et al.[1].

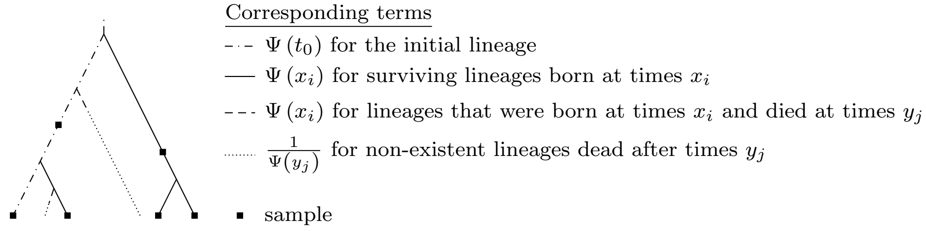


Figure S1: Complete tree integral mapping

##### S1.3 Reconstructed tree

In the complete tree all speciation, extinction and sampling events (through time and at present) are observed. However, in practice we are not able to observe (or infer) the complete tree, and therefore parameter estimation relies on the *reconstructed* tree. The reconstructed tree is a tree induced by the sampled nodes.

We define the function $E(t)$, the probability of a lineage alive at time $t$ to not leave sampled descendants. This function is described by the differential equation [1]:

$$\frac{\mathrm{d}E}{\mathrm{d}t} = -\left(\lambda + \mu + \psi\right) E + \lambda E^2 + \mu \tag{2}$$

with initial condition $E(0) = 1 - \rho_0$. Note that $1 - E(t_0)$ is the probability that at least one lineage was sampled either in the past or the present.

Denote

$$\Phi(t) = e^{\int_0^t 2\lambda(u)E(u) - \lambda(u) - \mu(u) - \psi(u)\, du}$$

Let $M_0$ be the number of extant species sampled. Then the probability density function of reconstructed trees conditioning on $(S)$, the event of sampling at least one lineage, is [1]:

$$\begin{aligned} & f^{\text{rec.}}\left(T \mid \lambda(t), \mu(t), \psi(t), \rho_0, \bar{t}, S\right) = \\ & \frac{\rho_0^{M_0}}{1 - E(t_0)} \Phi(t_0) \prod_{i=1}^{M_0+n-1} \lambda\left(x_i\right) \Phi(x_i) \\ & \times \prod_{j=1}^{n} \frac{\psi\left(y_j\right) E(y_j)}{\Phi(y_j)} \prod_{k=1}^{m} \psi\left(z_k\right) \end{aligned} \tag{3}$$

#### S2 Probability of not observing a lineage

The probability density function for the reconstructed trees (3) includes function $E$, which is defined as a solution to the differential equation (2). Therefore we can only consider the distribution of reconstructed trees when such a function exists and is unique. As we do not require rate functions to be continuous we consider solutions of the differential equation (2) in the extended sense [2, 3], that is, the equation for such solutions must hold almost everywhere on the interval $[0, t_0]$. We first formulate conditions for the rates that guarantee that any extended solution of equation (2) is bounded.

**Lemma S2.1.** Let $\lambda \geq 0$, $\mu \geq 0$, and $\psi \geq 0$ be Lebesgue-integrable on $[0, t_0]$, then any extended solution $E(t)$ of equation (2) will satisfy: $E(t) \geq 0$ on $[0, t_0]$.

*Proof.* Let $F(t) = \frac{E(t)}{\Psi(t)}$, taking the derivative of $F$ and using equation (2):

$$\frac{\mathrm{d}F}{\mathrm{d}t} = \lambda(t)\Psi(t)F^2 + \frac{\mu(t)}{\Psi(t)} \text{ for a.a. } t \in [0, t_0]$$

with $F(0) = 1 - \rho_0$. Note that the derivative of $F$ is non-negative for all $t \in [0, t_0]$ where it exists. Given that $F(0) \geq 0$ we have that $F(t) \geq 0$ for all $t \in [0, t_0]$. Therefore $E(t) \geq 0$ for all $t \in [0, t_0]$. □

**Lemma S2.2.** Let $\rho_0 > 0$,

- $\lambda \geq 0$, $\mu \geq 0$, and $\psi \geq 0$ are Lebesgue-integrable on $[0, t_0]$, and
- $\int_0^t \lambda(s) e^{-\int_0^s \lambda(u) - \mu(u) - \psi(u) du} ds < \infty$ for $t \in [0, t_0]$

then for any extended solution $E$: $E(t) < 1$ on $[0, t_0]$.

In the case when $\rho_0 = 0$, additional conditions:

- $\psi(0) > 0$
- $\lambda$, $\mu$, and $\psi$ are continuous in zero

guarantee $E(t) < 1$ on $(0, t_0]$.

*Proof.* Let $H(t) = [1 - E(t)]e^{-\int_0^t \lambda - \mu - \psi}$ then

$$\frac{\mathrm{d}H}{\mathrm{d}t} = -\lambda e^{\int_0^t (\lambda - \mu - \psi)} H^2 + \psi e^{-\int_0^t (\lambda - \mu - \psi)} \text{ for a.a. } t \in [0, t_0] \quad (4)$$

with $H(0) = \rho_0$. Note that $H(t) > 0$ implies $E(t) < 1$. We are going to prove that if $H(t^*) > 0$ for some $t^*$ then $H(t) > 0$ for all $t \in [t^*, t_0]$.

Suppose there is $\hat{t} \in (t^*, t_0]$ such that $H(\hat{t}) < 0$ then due to continuity of $H$ and $H(t^*) > 0$ there have to be $t' < t^*$ such that $H(t) > 0$ for all $t \in [t^*, t')$ and $H(t') = 0$. Then we have:

$$\frac{\mathrm{d}H}{\mathrm{d}t} \geq -\lambda e^{\int_0^t (\lambda - \mu - \psi)} H^2 \text{ for a. a. } t \in [t^*, t')$$

$$-\frac{\mathrm{d}\frac{1}{H}}{\mathrm{d}t} \geq -\lambda e^{\int_0^t (\lambda - \mu - \psi)} \text{ for a. a. } t \in [t^*, t')$$

$$\frac{\mathrm{d}\frac{1}{H}}{\mathrm{d}t} \leq \lambda e^{\int_0^t (\lambda - \mu - \psi)} \text{ for a. a. } t \in [t^*, t')$$

$$\frac{1}{H(t)} - \frac{1}{H(t^*)} \leq \int_{t^*}^{t} \lambda(s) e^{-\int_0^s \lambda(u)-\mu(u)-\psi(u)du} ds \text{ for a. a. } t \in [t^*, t')$$

$$H(t) \geq \frac{H(t^*)}{1 + H(t^*) \int_{t^*}^{t} \lambda(s) e^{-\int_0^s \lambda(u)-\mu(u)-\psi(u)du} ds} \text{ for a. a. } t \in [t^*, t')$$

$$H(t) \geq \frac{H(t^*)}{1 + H(t^*) \int_{t^*}^{t'} \lambda(s) e^{-\int_0^s \lambda(u)-\mu(u)-\psi(u)du} ds} = c > 0 \text{ for a. a. } t \in [t^*, t')$$

Due to the continuity of $H$ and $H(t') = 0$, there has to be a neighbourhood of $t'$ where $H$ is smaller than $c$ which contradicts the above inequality. Therefore $H(t) > 0$ for all $t \in [t^*, t_0]$.

Now if $\rho_0 > 0$, then $H(0) > 0$, and therefore $H(t) > 0$ for all $t \in [0, t_0]$. If $\rho_0 = 0$ and $\lambda$, $\mu$ and $\psi$ are continuous at zero, then $E$ is differentiable at zero and therefore $H$ is differentiable at zero. Furthermore due to $\psi(0) > 0$, the derivative of $H$ is strictly positive at zero. This implies that in the neighbourhood of zero $H$ increases and therefore $H(t) > 0$ in some neighbourhood of zero, which again implies that $H(t) > 0$ for all $t \in (0, t_0]$. □

We now show that the conditions of lemma S2.2 are sufficient for the existence and uniqueness of the solution to equation (2). Consider

$$f(t, E) = \lambda(t)E^2 - (\lambda(t) + \mu(t) + \psi(t))\, E + \mu(t)$$

on $[0, t_0] \times [0, 1]$. $f$ is Lebesgue-integrable in $t$ for any fixed $E$ and continuous in $E$ for any fixed $t$. From $0 \leq E(t) \leq 1$ we have:

$$|f(t, E)| \leq \lambda(t)E^2 + (\lambda(t) + \mu(t) + \psi(t))\, E + \mu(t) \leq$$

$$2\lambda(t) + 2\mu(t) + \psi(t) \text{ for } (t, E) \in [0, t_0] \times [0, 1]$$

As the right-hand side of the inequality is Lebesgue-integrable, $f$ satisfies the conditions of the Carathéodory's existence theorem [2, 3] and therefore there is a local solution (in the extended sense) to equation (2), which can be extended on the whole interval $[0, t_0]$ due to $E$ being bounded [2, 3].

$f$ is locally lipschitzian on $[0, t_0] \times [0, 1]$:

$$|f(t, E_1) - f(t, E_2)| = \Big| \big[\lambda(t)(E_1 + E_2) - (\lambda(t) + \mu(t) + \psi(t))\big](E_1 - E_2) \Big| \leq$$

$$(3\lambda(t) + \mu(t) + \psi(t))|E_1 - E_2|$$

for any $t \in [0, t_0]$ with $3\lambda(t) + \mu(t) + \psi(t)$ being Lebesgue-integrable. Therefore the solution is unique [2, 3].

Note that the classic existence and uniqueness theorems used here require the domain of $f$ to be open. To achieve this one can extend $f$ to $(-\epsilon, t_0+\epsilon)\times(-\epsilon, 1+\epsilon)$ for some $\epsilon > 0$ by setting $f$ to zero whenever $t$ is outside of $[0, t_0]$ and setting $f(t, E) = f(t, 1)$ for $E > 1$ and $f(t, E) = f(t, 0)$ for $E < 0$. Then the above requirements of existence and uniqueness will still hold for the extended domain and given that any solution with the initial condition in $0 \times [0, 1]$ will not escape

$[0, t_0] \times [0, 1]$, the maximal interval of existence for such solutions will contain $[0, t_0]$.

The conditions of lemma S2.2 guarantee that we can define a probability distribution over the reconstructed trees. We could not relax these conditions and did not attempt to show that they are necessary conditions. Milder conditions could possibly be found.

### S3 Elements of complete FBD identifiability proof

#### S3.1 Probability space on trees

We first define a measure on the space of trees. Consider a complete tree $\mathcal{T}$ that is obtained under the time-dependent birth-death model without present sampling (guaranteed complete sampling) or sampling through time. Its discrete structure is represented by a ranked oriented topology [4]. Ranked means that each branching or tip node has a distinct ranking, except the extant tips, which all have rank $N_0 + n$, and the ranking is consistent with the ranking induced by ancestor-descendant relationship in the tree. Oriented means that every branch after a branching event is designated by a label *left* or *right*. Each ranked node is mapped to a time point on the interval $[0, t_0]$ preserving the ranking. Thus, $\mathcal{T} = (T, t_1, \dots, t_{N_0+n-1})$, where a node with rank $i$ has time $t_i$. In the notation introduced above, $t_i = x_{\pi_x(i)}$ and $t_i = y_{\pi_y(i)}$, with corresponding functions $\pi_x$ and $\pi_y$ that map indices of branching and death events to the rankings on the tree. Thus, we can see the space of trees as a disjoint union of sets $S_T = \{(t_1, \dots, t_{N_0+n-1}) \mid t_0 > t_1 > \dots > t_{N_0+n-1} > 0\}$ for each distinct ranked topology $T$. Then the probability measure of a set $\mathbb{T}$ of trees is $\mu(\mathbb{T}) = \sum_{T \in \mathbf{T}} \mu(S_T \bigcap \mathbb{T})$ where $\mathbf{T}$ is the set of all ranked topologies.

It is straightforward to extend this definition to the case with sampling through time and present sampling. In the case of sampling through time, the topology will have extra nodes representing sampling events with corresponding rankings and the vector of times will have additional elements for the times of the sampled nodes. Nodes sampled at present will just be designated nodes in the topology component.

#### S3.2 Parameter constraints for equal probability densities

The identifiability of the time-dependent FBD model for complete trees relies on Lemma 0.3 from the main text. Here we provide a more detailed proof of the lemma.

*Proof.* Remember that $t_0$ is fixed, $\theta_1 = (\lambda_1, \mu_1, \psi_1, t_0)$, and $\theta_2 = (\lambda_2, \mu_2, \psi_2, t_0)$. Assume that for all ranked topologies $T$,

$$f^{\text{compl.}}(T, \bar{t}|\theta_1) = f^{\text{compl.}}(T, \bar{t}|\theta_2)$$

for all $\bar{t} \in S_T \setminus \mathbf{0}_T$ where $\boldsymbol{\mu}(\mathbf{0}_T) = 0$.

Consider all trees with a single birth event and no death or sampling events. An example of such a tree is given in Figure S2a. All such trees will have the same ranked topology $G$ and only differ by the time of the branching event $x_1$. For each such tree $(G, x_1)$ we define a tree which is produced by adding one extant lineage at a time point $x_a$ after the first birth event on the right branch, e.g., $x_a < x_1$ (Figures S2b). Denote the obtained tree $(G^+, x_1, x_a)$. We also define a tree $(G^*, x_1, y_a)$ (Figures S2c) formed by contracting the rigth extant edge in $(G, x_1)$ to become extinct at time $y_a$ after the birth event.

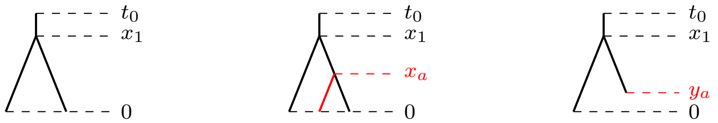


(a) $G$ (b) $G^+$ (c) $G^*$

Figure S2: Tree variants. Subfigure (a) depicts the original tree $G$, while Subfigure (b) shows the addition of an extant edge at time $x_a$ and Subfigure (c) shows the contraction of an extant edge such that it dies at time $y_a$.

Note $S_G = [0, t_0]$ and $S_{G^+} = \{(x_1, x_a) \mid t_0 > x_1 > x_a > 0\}$. Since the probability densities for model 1 with parameters $\theta_1$ and model 2 with parameters $\theta_2$ are equal almost everywhere, we have:

$$f^{\text{compl.}}(G, x_1|\theta_1) = f^{\text{compl.}}(G, x_1|\theta_2)$$

for all $x_1 \in S_G \setminus \mathbf{0}_G$ where $\boldsymbol{\mu}(\mathbf{0}_G) = 0$.

We also have:

$$f^{\text{compl.}}(G^+, x_1, x_a|\theta_1) = f^{\text{compl.}}(G^+, x_1, x_a|\theta_2)$$

for all $(x_1, x_a) \in S_{G^+} \setminus \mathbf{0}_{G^+}$ and $\boldsymbol{\mu}(\mathbf{0}_{G^+}) = 0$.

Using equation (1) for the probability density of complete trees, we can rewrite the last two equations as follows:

$$\Psi_1(t_0)\lambda_1(x_1)\Psi_1(x_1) = \Psi_2(t_0)\lambda_2(x_1)\Psi_2(x_1)$$
$$\Psi_1(t_0)\lambda_1(x_1)\Psi_1(x_1)\lambda_1(x_a)\Psi_1(x_a) = \Psi_2(t_0)\lambda_2(x_1)\Psi_2(x_1)\lambda_2(x_a)\Psi_2(x_a)$$

Therefore

$$\lambda_1(x_a)\,\Psi_1(x_a) = \lambda_2(x_a)\,\Psi_2(x_a) \text{ for all } x_a \in [0, t_0] \setminus \mathbf{0}_1 \tag{5}$$

where

$$\mathbf{0}_1 = \{x \mid \text{ for all } x' \in (x, t_0] \setminus \mathbf{0}_G : (x', x) \in \mathbf{0}_{G^+}\}$$

Now we show that $\boldsymbol{\mu}(\mathbf{0}_1) = 0$. Suppose $\boldsymbol{\mu}(\mathbf{0}_1) > 0$ then for some $b < t_0$: $\boldsymbol{\mu}(\mathbf{0}_1 \cap [0, b]) > 0$. The product of non-zero measure sets is non-zero: $\boldsymbol{\mu}((\mathbf{0}_1 \cap [0, b]) \times ((b, t_0] \setminus \mathbf{0}_G)) > 0$. Note that $(\mathbf{0}_1 \cap [0, b]) \times ((b, t_0] \setminus \mathbf{0}_G) \subset \mathbf{0}_{G^+}$, and therefore $\boldsymbol{\mu}(\mathbf{0}_{G^+}) > 0$, which contradicts the definition of $\mathbf{0}_{G^+}$, therefore $\boldsymbol{\mu}(\mathbf{0}_1) = 0$.

Applying a similar argument to trees $G$ and $G^*$ we obtain:

$$\frac{\mu_1\left(y_a\right)}{\Psi_1(y_a)} = \frac{\mu_2\left(y_a\right)}{\Psi_2(y_a)} \text{ for all } y_a \in [0, t_0] \setminus \mathbf{0}_2 \tag{6}$$

Then (5) and (6) imply:

$$\lambda_1(t)\mu_1(t) = \lambda_2(t)\mu_2(t) \text{ for all } t \in [0, t_0] \setminus (\mathbf{0}_1 \cup \mathbf{0}_2) \tag{7}$$

A similar argument applied to trees where one sampling event is added after the birth event on one of the branches would lead to $\psi_1(t) = \psi_2(t)$ for all $t \in [0, t] \setminus \mathbf{0}_3$ with $\boldsymbol{\mu}(\mathbf{0}_3) = 0$.

□

#### S3.3 Uniqueness of gamma

Denote $\gamma(t) = \frac{\Psi_1(t)}{\Psi_2(t)}$. Note that due to $\lambda_1$, $\lambda_2$, $\mu_1$, $\mu_2$, $\psi_1$ and $\psi_2$ being Lebesgue-integrable on $[0, t_0]$, $\gamma(t)$ is continuous on the interval $[0, t_0]$ and therefore it is bounded by some constant $K$ on this interval.

The parameter constraints (see Lemma (0.3) in the main text) imply that:

$$\begin{aligned}
\lambda_2\left(t\right) &= \gamma\left(t\right)\lambda_1\left(t\right) \text{ for a. a. } t \in [0, t_0] \\
\mu_2\left(t\right) &= \frac{\mu_1\left(t\right)}{\gamma\left(t\right)} \text{ for a. a. } t \in [0, t_0] \\
\psi_2(t) &= \psi_1(t) \text{ for a. a. } t \in [0, t_0]
\end{aligned}$$

Using these equations and the definition of $\gamma$ we obtain:

$$\gamma\left(t\right) = e^{-\int_0^t \lambda_1 + \mu_1 - \gamma\lambda_1 - \frac{\mu_1}{\gamma}\, du} \text{ for a. a. } t \in [0, t_0]$$

Taking the derivative of both sides we have:

$$\frac{d\gamma}{dt} = \lambda_1\gamma^2 - (\lambda_1 + \mu_1)\gamma + \mu_1 \text{ for a. a. } t \in [0, t_0]$$

From the definition of $\gamma$ we obtain the initial condition $\gamma(0) = 1$. Moreover, from Equation (0.8) in the main text we have that $\gamma(t_0) = 1$. An extended solution to this equation is $\gamma(t) = 1$.

The following inequality holds for the right hand side of the equation:

$$|\lambda_1\gamma_1^2 - (\lambda_1+\mu_1)\gamma_1 + \mu_1 - (\lambda_1\gamma_2^2 - (\lambda_1+\mu_1)\gamma_2 + \mu_1)| \leq |\gamma_1 + \gamma_2 - \lambda_1 - \mu_1||\gamma_1 - \gamma_2| \leq$$

$$(2K_U + \lambda_1 + \mu_1)|\gamma_1 - \gamma_2|$$

where $K = sup_{t \in U}\gamma(t)$ for a given compact set $U$. Since $2K + \lambda_1 + \mu_1$ is Lebesgue-integrable the solution is unique [2, 3].

### S4 Transformed (pulled) rates

In the main text, we introduced a parameter transformation (repeated from equation (0.10) in the main text) that defines pulled rates:

$$\begin{aligned}\tilde{\lambda} &= \lambda(1-E)\\ \tilde{\mu} &= \frac{\psi E}{1-E}\\ \tilde{\psi} &= \psi\end{aligned} \tag{8}$$

Note that in the conditions of lemma S2.2, $E(t) < 1$ and $\tilde{\mu}$ is defined for all $t \in (0, t_0]$.

#### S4.1 Reconstructed and complete tree distributions

In this section, we show that the distribution of reconstructed trees is equivalent to the distribution of complete trees generated by the FBD process with transformed parameters.

Let $f^{\text{compl.}}(\mathcal{T}|\tilde{\lambda}, \tilde{\mu}, \tilde{\psi}, t_0)$ be the probability density of complete trees under the time-dependent FBD model with $\tilde{\rho}_0 = 1$. Let $f^{\text{rec.}}(\mathcal{T}|\lambda, \mu, \psi, \rho_0, t_0, S)$ be the probability density of reconstructed trees under the time-dependent FBD model, conditioned on sampling of at least one lineage ($S$).

The probability density of the complete trees under the pulled rates is (in what follows we omit $t_0$ to simplify notation):

$$\begin{aligned}f^{\text{compl.}}\left(\mathcal{T} \mid \tilde{\lambda}, \tilde{\mu}, \tilde{\psi}\right) =& e^{-\int_0^{t_0} \tilde{\lambda}(u)+\tilde{\mu}(u)+\tilde{\psi}(u)\,du}\\ &\times \prod_{i=1}^{M_0+n-1} \tilde{\lambda}\left(x_i\right) e^{-\int_0^{x_i} \tilde{\lambda}(u)+\tilde{\mu}(u)+\tilde{\psi}(u)\,du}\\ &\times \prod_{j=1}^{n} \tilde{\mu}\left(y_j\right) e^{\int_0^{y_j} \tilde{\lambda}(u)+\tilde{\mu}(u)+\tilde{\psi}(u)\,du} \prod_{k=1}^{m} \tilde{\psi}\left(z_k\right)\end{aligned}$$

Substituting the values for the pulled rates in the exponential terms we obtain:

$$\begin{aligned}e^{-\int_0^t \tilde{\lambda}+\tilde{\mu}+\tilde{\psi}\,dt} =& e^{-\int_0^t \lambda-\lambda E+\frac{\psi E}{1-E}+\psi\,dt}\\ =& e^{-\int_0^t \lambda E-\mu+\frac{\psi E}{1-E}\,dt}\left(e^{-\int_0^t \lambda+\mu+\psi-2\lambda E\,dt}\right)\end{aligned}$$

Assuming $\rho_0 > 0$ and using equation (2), we have:

$$\begin{aligned}e^{\int_0^t \lambda E-\mu+\frac{\psi E}{1-E}\,dt} =& e^{\int_0^t \frac{\frac{dE}{dt}}{E-1}\,du} = e^{\ln|E(t)-1|-\ln|E(0)-1|}\\ =& e^{\ln\frac{1-E(t)}{\rho_0}} = \frac{1-E\left(t\right)}{\rho_0}\end{aligned}$$

which implies:

$$e^{-\int_0^t \tilde{\lambda}+\tilde{\mu}+\tilde{\psi}\,dt} = \frac{\rho_0}{1-E(t)}\left(e^{-\int_0^t \lambda+\mu+\psi-2\lambda E\,dt}\right)$$

We thus re-write the complete tree probability density using exponential terms of the same form as the reconstructed tree probability density.

$$\begin{aligned} f^{\text{compl.}}\left(\mathcal{T} \mid \tilde{\lambda}, \tilde{\mu}, \tilde{\psi}\right) =& \frac{\rho_0}{1-E(t_0)} e^{-\int_0^{t_0} \lambda+\mu+\psi-2\lambda E\,du} \\ &\times \prod_{i=1}^{M_0+n-1} \tilde{\lambda}(x_i) \frac{\rho_0}{1-E(x_i)} e^{-\int_0^{x_i} \lambda+\mu+\psi-2\lambda E\,du} \\ &\times \prod_{j=1}^{n} \tilde{\mu}(y_j) \frac{(1-E(y_j))}{\rho_0} e^{\int_0^{y_j} \lambda+\mu+\psi-2\lambda E\,du} \prod_{k=1}^{m} \tilde{\psi}(z_k) \end{aligned}$$

We substitute the pulled rates and use Equation (0.3) in the main article:

$$\begin{aligned} & f^{\text{compl.}}\left(\mathcal{T} \mid \tilde{\lambda}, \tilde{\mu}, \tilde{\psi}\right) = \\ & \frac{\rho_0}{1-E(t_0)} e^{-\int_0^{t_0} \lambda+\mu+\psi-2\lambda E\,du} \\ & \times \prod_{i=1}^{M_0+n-1} \lambda(x_i)(1-E(x_i)) \frac{\rho_0}{1-E(x_i)} e^{-\int_0^{x_i} \lambda+\mu+\psi-2\lambda E\,du} \\ & \times \prod_{j=1}^{n} \left(\psi(y_j) \frac{E(y_j)}{\rho_0}\right) e^{\int_0^{y_j} \lambda+\mu+\psi-2\lambda E\,du} \prod_{k=1}^{m} \psi(z_k) \\ & = f^{\text{rec.}}(\mathcal{T} \mid \lambda, \mu, \psi, S) \end{aligned}$$

When $\rho_0 = 0$, similar derivations imply $e^{\int_y^x \lambda E-\mu+\frac{\psi E}{1-E}\,dt} = \frac{1-E(x)}{1-E(y)}$. Without present day samples, each branching event can be matched with a death event and exponential terms in both complete and reconstructed densities can be written as: $e^{-\int_{y_j}^{x_i} \tilde{\lambda}+\tilde{\mu}+\tilde{\psi}\,du}$ or $e^{-\int_{y_j}^{x_i} \lambda+\mu+\psi-2\lambda E\,du}$. Then substitution of pulled rates in the complete tree density will again produce the reconstructed tree density conditioned on sampling of at least one lineage. Note that in this border case, $E(0) = 1$ and the pulled birth rate becomes zero while pulled death rate approaches infinity as time approaches zero. This guarantees there are no lineages surviving until present in the complete tree case.

#### S4.2 Connection to deterministic FBD process

A deterministic birth-death sampling model provides an equation for the lineage through time (LTT) curve based on the birth, death, and sampling rates. This LTT curve corresponds to the expected number of lineages through time in the stochastic birth-death sampling model [5] with the same rates and some sampling fraction. Considering the deterministic approach, the pulled rates describe the expected number of observed lineages (occurring in the reconstructed tree) in the corresponding stochastic model. In what follows we derive the pulled rates using the deterministic approach.

##### Probability of unobserved lineages

The pulled birth rate $\tilde{\lambda} = \lambda (1 - E)$ has been derived in Louca and Pennell's unidentifiability result for reconstructed timetrees. This expression falls naturally from the definition of $E(t)$ as "the fraction of lineages extant at time $t$ that will be missing from the timetree (either due to extinction or not having been sampled)" [6]. Then, only the birth events that produce surviving lineages will contribute to the pulled birth rate, hence the expression for this rate.

The differential equation (2) that describes $E$ is not guaranteed to have an analytical solution, and therefore it is useful to also view $E(t)$ from an alternative perspective. Namely, $1 - E(t)$ is also the percentage of lineages alive at time $t$ that we will find in the sample under the deterministic model based on birth, death and sampling rates. It is easy to define a pulled death rate similarly to the pulled birth rate, and produce an expression describing the number of observed lineages alive at time $t$, which allows the proportion of unobserved lineages to be calculated.

From Louca and Pennell [6], we have the expected number of lineages, both observed and unobserved, denoted $N(t)$. $M_0$ is the number of lineages sampled at the present and $\rho_0$ is the sampling fraction.

$$N(t) = \frac{M_0}{\rho_0} e^{\int_0^t \mu(u) - \lambda(u)\, du}$$

Let $V(t)$ be the expected number of observed lineages. Note that the number of observed lineages under the original model is equal to the number of all lineages in the complete trees under the model with pulled rates:

$$V(t) = M_0 e^{\int_0^t \tilde{\mu}(u) - \tilde{\lambda}(u)\, du}$$

The probability that a lineage will be missing from the reconstructed timetree is equal to the proportion of lineages at that time that are unobserved.

$$\begin{aligned} E(t) =& \frac{N(t) - V(t)}{N(t)} \\ =& 1 - \rho_0 e^{\int_0^t \tilde{\mu}(u) - \tilde{\lambda}(u) - \mu(u) + \lambda(u)\, du} \end{aligned} \tag{9}$$

##### Pulled death rate

From equation (9) and the expression for the pulled birth rate, we obtain:

$$E(t) = 1 - \rho_0 e^{\int_0^t \tilde{\mu}(u) + \lambda(u) E(u) - \mu(u)\, du}$$

By calculating the derivative of both sides and combining it with Equation (2), we find

$$\tilde{\mu} + \lambda E - \mu = \lambda E - \mu - \frac{\psi E}{E - 1}$$

which implies

$$\tilde{\mu} = \frac{\psi E}{1 - E}$$

###### Pulled sampling rate

As the sampling rate on observed lineages is the same as in the initial process, we set $\tilde{\psi} = \psi$.

#### S4.3 One-to-one correspondence between pulled and normal rates

Now we show that Equation (8) defines a one-to-one correspondence between sets of birth, death, and strictly positive sampling rates and corresponding pulled rates.

We say that two sets of rates $F_1 = (\lambda_1, \mu_1, \psi_1, \rho_{0,1})$ and $F_2 = (\lambda_2, \mu_2, \psi_2, \rho_{0,2})$ are not equal ($F_1 \neq F_2$) if there is a pair of rates (birth, death, or sampling) that are not equal everywhere on $[0, t_0]$ or $\rho_{0,1} \neq \rho_{0,2}$. We also write $\alpha_1 \overset{\text{a.e.}}{=} \alpha_2$ for functions that are equal almost everywhere on $[0, t_0]$ and $\alpha_1 \overset{\text{a.e.}}{\neq} \alpha_2$ for functions that differ on at least a non-zero measure subset of $[0, t_0]$.

**Lemma S4.1.** Let $F_1 = (\lambda_1, \mu_1, \psi_1, \rho_{0,1})$ and $F_2 = (\lambda_2, \mu_2, \psi_2, \rho_{0,2})$ and $F_1 \neq F_2$. Let $\tilde{F}_1 = \left(\tilde{\lambda}_1, \tilde{\mu}_1, \tilde{\psi}_1\right)$ and $\tilde{F}_2 = \left(\tilde{\lambda}_2, \tilde{\mu}_2, \tilde{\psi}_2\right)$ be the pulled rates obtained from $F_1$ and $F_2$ using Equation (8). Then $\tilde{F}_1 \neq \tilde{F}_2$.

*Proof.* We consider two cases.

**Case 1**: $\psi_1 \overset{\text{a.e.}}{\neq} \psi_2$

The case where $\psi_1 \overset{\text{a.e.}}{\neq} \psi_2$ is trivial, as we know $\tilde{\psi}_1 \overset{\text{a.e.}}{=} \psi_1$ and $\tilde{\psi}_2 \overset{\text{a.e.}}{=} \psi_2$, implying $\tilde{\psi}_1 \overset{\text{a.e.}}{\neq} \tilde{\psi}_2$ and therefore $\tilde{F}_1 \neq \tilde{F}_2$.

**Case 2**: $\psi_1 \overset{\text{a.e.}}{=} \psi_2$

We now consider the case where $\psi_1 \overset{\text{a.e.}}{=} \psi_2$. Suppose $\tilde{F}_1 = \tilde{F}_2$. Then from $\tilde{\mu}_1 \overset{\text{a.e.}}{=} \tilde{\mu}_2$:

$$\frac{\psi_1 E_1}{1 - E_1} \overset{\text{a.e.}}{=} \frac{\psi_2 E_2}{1 - E_2}$$

As we assume that $\psi_1(t), \psi_2(t) > 0$ for almost all $t \in [0, t_0]$, we have $E_1(t), E_2(t) < 1$ for all $t$ except possibly for $t = 0$ in the case when $\rho_0 = 0$.

$$\frac{1-E_1}{E_1} \overset{\text{a.e.}}{=} \frac{1-E_2}{E_2}$$
$$\frac{1}{E_1} - 1 \overset{\text{a.e.}}{=} \frac{1}{E_2} - 1$$
$$E_1 \overset{\text{a.e.}}{=} E_2$$

From continuity of $E_1$ and $E_2$ we have $E_1 = E_2$ for all $t \in [0, 1]$ and therefore $\rho_{0,1} = 1 - E_1(0) = 1 - E_2(0) = \rho_{0,2}$.

From $\tilde{\lambda}_1 = \tilde{\lambda}_2$:

$$\lambda_1 (1 - E_1) \overset{\text{a.e.}}{=} \lambda_2 (1 - E_2)$$
$$\lambda_1 (1 - E_1) \overset{\text{a.e.}}{=} \lambda_2 (1 - E_1)$$
$$\lambda_1 \overset{\text{a.e.}}{=} \lambda_2$$

From $E_1 = E_2$, we know that their derivatives exist almost everywhere and are also equal when they exist:

$$\frac{\mathrm{d}E_1}{\mathrm{d}t} \overset{\text{a.e.}}{=} \frac{\mathrm{d}E_2}{\mathrm{d}t}$$
$$-(\lambda_1 + \mu_1 + \psi_1) E_1 + \lambda_1 E_1^2 + \mu_1 \overset{\text{a.e.}}{=} -(\lambda_2 + \mu_2 + \psi_2) E_2 + \lambda_2 E_2^2 + \mu_2$$
$$-(\lambda_1 + \mu_1 + \psi_1) E_1 + \lambda_1 E_1^2 + \mu_1 \overset{\text{a.e.}}{=} -(\lambda_1 + \mu_2 + \psi_1) E_1 + \lambda E_1^2 + \mu_2$$
$$\mu_1 (1 - E_1) \overset{\text{a.e.}}{=} \mu_2 (1 - E_1)$$
$$\mu_1 \overset{\text{a.e.}}{=} \mu_2$$

This contradicts the assumption of $F_1 \neq F_2$, therefore $\tilde{F}_1 \neq \tilde{F}_2$ □

#### S4.4 Reverse transformation

Equation (8) (also equation (0.10) in the main text) defines the transformation of the FBD process rates to pulled rates. Here we derive the reverse transformation, where rates of the process producing reconstructed trees are defined through pulled rates (rates of the process that produces complete trees with the same distribution). Assume the pulled rates are non-negative for all $t$ and $\tilde{\psi}(0) > 0$ for a. a. $t$. We also assume that $\mu$ and $\psi$ are piecewise differentiable. Then the reverse transformation is:

$$
\begin{aligned}
\lambda &= \frac{\tilde{\lambda}(\tilde{\mu}+\tilde{\psi})}{\tilde{\psi}} \\
\mu &= \frac{\mathrm{d}\frac{\tilde{\mu}}{\tilde{\psi}}}{\mathrm{d}t}\frac{\tilde{\psi}}{\tilde{\mu}+\tilde{\psi}} + \tilde{\mu}\frac{\tilde{\lambda}+\tilde{\psi}}{\tilde{\psi}} \\
\psi &= \tilde{\psi} \\
\rho_0 &= \frac{\tilde{\psi}(0)}{\tilde{\psi}(0)+\tilde{\mu}(0)}
\end{aligned}
\tag{10}
$$

Now we show that this is indeed the reverse transformation of the pulled rates into original rates. Firstly, $\tilde{\psi}$ determines $\psi$. To determine $\tilde{\lambda}$, we can write $E$ in terms of the pulled death and sampling rates:

$$
\begin{aligned}
E =& \frac{\tilde{\mu}}{\tilde{\psi}}(1-E) \\
E =& \frac{\frac{\tilde{\mu}}{\tilde{\psi}}}{1+\frac{\tilde{\mu}}{\tilde{\psi}}}
\end{aligned}
$$

which gives us:

$$
E = \frac{\tilde{\mu}}{\tilde{\mu}+\tilde{\psi}} \tag{11}
$$

Having the expression for $E$ in terms of the pulled rates the expression for $\lambda$ is

$$
\lambda = \frac{\tilde{\lambda}\left(\tilde{\mu}+\tilde{\psi}\right)}{\tilde{\psi}}
$$

Furthermore as $\rho_0 = 1 - E(0)$, we obtain

$$
\rho_0 = \frac{\tilde{\psi}(0)}{\tilde{\psi}(0)+\tilde{\mu}(0)}
$$

To determine $\mu$, we first write $\frac{\mathrm{d}E}{\mathrm{d}t}$ with respect to the pulled death and sampling rates.

$$\begin{aligned}\frac{\mathrm{d}E}{\mathrm{d}t} =& \frac{\mathrm{d}\frac{\tilde{\mu}}{\tilde{\mu}+\tilde{\psi}}}{\mathrm{d}t}\\ =& \frac{\frac{\mathrm{d}\tilde{\mu}}{\mathrm{d}t}\left(\tilde{\mu}+\tilde{\psi}\right)-\tilde{\mu}\left(\frac{\mathrm{d}\tilde{\mu}}{\mathrm{d}t}+\frac{\mathrm{d}\tilde{\psi}}{\mathrm{d}t}\right)}{\left(\tilde{\mu}+\tilde{\psi}\right)^2}\\ =& \frac{\frac{\mathrm{d}\tilde{\mu}}{\mathrm{d}t}\tilde{\psi}-\tilde{\mu}\frac{\mathrm{d}\tilde{\psi}}{\mathrm{d}t}}{\left(\tilde{\mu}+\tilde{\psi}\right)^2}\\ =& \frac{\tilde{\psi}^2}{\left(\tilde{\mu}+\tilde{\psi}\right)^2}\frac{\mathrm{d}\frac{\tilde{\mu}}{\tilde{\psi}}}{\mathrm{d}t}\end{aligned}$$

From equation (2) and $\tilde{\lambda}+\tilde{\psi}=\lambda(1-E)+\psi$ we have:

$$\begin{aligned}\mu\left(1-E\right) =& \frac{\mathrm{d}E}{\mathrm{d}t}+(\tilde{\lambda}+\tilde{\psi})E\\ \frac{\tilde{\psi}}{\tilde{\mu}+\tilde{\psi}}\mu =& \frac{\tilde{\psi}^2}{\left(\tilde{\mu}+\tilde{\psi}\right)^2}\frac{\mathrm{d}\frac{\tilde{\mu}}{\tilde{\psi}}}{\mathrm{d}t}+\left(\tilde{\lambda}+\tilde{\psi}\right)\frac{\tilde{\mu}}{\tilde{\mu}+\tilde{\psi}}\\ \mu =& \frac{\tilde{\psi}}{\tilde{\mu}+\tilde{\psi}}\frac{\mathrm{d}\frac{\tilde{\mu}}{\tilde{\psi}}}{\mathrm{d}t}+\left(\tilde{\lambda}+\tilde{\psi}\right)\frac{\tilde{\mu}}{\tilde{\psi}}\end{aligned}$$

### S5 Adding removal at sampling probability

Here we consider the time-dependent birth-death sampling model with an extra parameter, removal at sampling probability $r$. In such a process a lineage sampled at time $t$ is removed from the process with probability $r\left(t\right)$. In the main text we showed that if $r$ is an arbitrary time-dependent function then the time-dependent birth-death sampling model with removal at sampling is unidentifiable. Here we show that when $r$ is a fixed function and $r(t)<1$ for almost all $t\in[0,t_0]$, the parameter transformation (equation 10 from the main text) again defines a one-to-one correspondence between the distributions of reconstructed trees generated by time-dependent birth-death sampling models with removal at sampling and distributions of complete trees generated by the FBD model with pulled rates.

#### S5.1 Pulled rates

The pulled rates for the reconstructed tree are as follows. Note that the pulled birth rate does not differ from the $r\left(t\right)=0$ case.

$$\begin{aligned}\tilde{\lambda} &= \lambda(1-E)\\ \tilde{\mu} &= \psi\left(\frac{E}{1-E}+r\right)\\ \tilde{\psi} &= \psi(1-r)\end{aligned} \tag{12}$$

The probability $E(t)$ of a lineage alive at time $t$ to not leave sampled descendants is independent of what happens to lineages after being sampled. Therefore $E$ and differential equation (2) remain **unchanged**.

$N(t)$, the total number of lineages both observed and unobserved, is affected by $r(t)$, while $V(t)$, the total number of observed lineages, is not affected.

$$N(t) = \frac{M_0}{\rho_0} e^{\int_0^t \mu+\psi r-\lambda\, dt}$$

$$V(t) = M_0 e^{\int_0^t \tilde{\mu}-\tilde{\lambda}\, dt}$$

Therefore, we obtain a new expression for $\frac{\mathrm{d}E}{\mathrm{d}t}$:

$$\begin{aligned}E(t) =& \frac{N(t)-V(t)}{N(t)}\\ =& 1-\rho_0 e^{\int_0^t \tilde{\mu}+\lambda E-\mu-\psi r\, dt}\\ \frac{\mathrm{d}E}{\mathrm{d}t} =& (E-1)(\tilde{\mu}+\lambda E-\mu-\psi r)\end{aligned}$$

We can use our two expressions for $\frac{\mathrm{d}E}{\mathrm{d}t}$ to determine the pulled death rate.

$$\tilde{\mu}+\lambda E-\mu-\psi r = \lambda E-\mu+\frac{\psi E}{1-E}$$

$$\tilde{\mu} = \frac{\psi E}{1-E}+\psi r$$

#### S5.2 Probability density equality

MacPherson et al. [1] gives the probability density function for reconstructed trees generated by the birth-death sampling process with removal at sampling, conditioned on S, the event of sampling at least one lineage, as:

$$\begin{aligned}f^{\text{rec.}}(\mathcal{T}|\lambda,\mu,\psi,r,t_0,S) =& \frac{\rho_0^{M_0}}{1-E(t_0)}\Phi(t_0)\prod_{k=1}^{m}\psi(z_k)(1-r(z_k))\\ &\times \prod_{i=1}^{M_0+n-1}\lambda(x_i)\Phi(x_i)\\ &\times \prod_{j=1}^{n}\frac{\psi(y_j)}{\Phi(y_j)}((1-r(y_j))E(y_j)+r(y_j))\end{aligned}$$

To show that this distribution is the same as the distribution of complete trees under the FBD model with pulled rates we again substitute pulled rates in the expression for the probability density $f^{\text{compl.}}\left(\mathcal{T}|\tilde{\lambda},\tilde{\mu},\tilde{\psi},t_0\right)$ (equation 1) with expressions for the pulled rates defined by equations (12). The contribution of speciation events in these densities remains unchanged compared to the case when $r(t)=0$. The equality of sampling event terms is obvious and it only remains to consider death events and affected edges to show that the complete and reconstructed tree probability densities remain equal.

We confirm the equality connecting the exponential terms previously shown for $r(t)=0$:

$$\begin{aligned}
e^{-\int_0^t \tilde{\lambda}+\tilde{\mu}+\tilde{\psi}\,dt} =& e^{-\int_0^t \lambda-\lambda E+\frac{\psi E}{1-E}+\psi r+\psi(1-r)\,dt} \\
=& \left(e^{-\int_0^t -\mu+\lambda E+\frac{\psi E}{1-E}\,dt}\right)e^{-\int_0^t \lambda+\mu+\psi-2\lambda E\,dt} \\
=& \frac{\rho_0}{1-E}e^{-\int_0^t \lambda+\mu+\psi-2\lambda E\,dt}
\end{aligned}$$

$$\begin{aligned}
\tilde{\mu}e^{\int_0^t \tilde{\lambda}+\tilde{\mu}+\tilde{\psi}\,dt} =& \left(\frac{\psi E}{1-E}+\psi r\right)\left(\frac{1-E}{\rho_0}\right)\frac{1}{\Phi(t)} \\
=& \frac{\psi}{\rho_0}\left(E+r-rE\right)\frac{1}{\Phi(t)} \\
=& \frac{\psi}{\rho_0\Phi(t)}\left((1-r)E+r\right)
\end{aligned}$$

Thus, the distributions are equal.

#### S5.3 One-to-one correspondence between pulled and normal rates

**Lemma S5.1.** Let $F_1=(\lambda_1,\mu_1,\psi_1,\rho_{0,1},r)$ and $F_2=(\lambda_2,\mu_2,\psi_2,\rho_{0,2},r)$ where $r$ is a fixed function such that $r(t)<1$ for almost all $t\in[0,t_0]$, and $F_1\neq F_2$. Let $\tilde{F}_1=\left(\tilde{\lambda}_1,\tilde{\mu}_1,\tilde{\psi}_1\right)$ and $\tilde{F}_2=\left(\tilde{\lambda}_2,\tilde{\mu}_2,\tilde{\psi}_2\right)$ be the pulled rates obtained from $F_1$ and $F_2$ using Equation (8). Then $\tilde{F}_1\neq\tilde{F}_2$.

The proof of Lemma S5.1 closely follows the proof of Lemma S4.1. This lemma implies the identifiability of the time-dependent birth-death-sampling model with fixed removal probability.

#### S5.4 Original rates from pulled rates

As previously, we derive expression for $(E,\lambda,\mu,\psi,\rho_0)$ with respect to the pulled rates and $r$:

$$
\begin{aligned}
\lambda &= \frac{\tilde{\lambda}\left(1-r\right)\left(\tilde{\psi}+\tilde{\mu}\right)}{\tilde{\psi}} \\
\mu &= \tilde{\mu}\left(\frac{\tilde{\psi}+\tilde{\lambda}-\tilde{\lambda}r}{\tilde{\psi}}\right)-\tilde{\lambda}r-\frac{\tilde{\psi}r}{1-r}+\frac{\mathrm{d}\frac{\tilde{\mu}}{\tilde{\psi}}}{\mathrm{d}t}\frac{\tilde{\psi}}{\tilde{\mu}+\tilde{\psi}}-\frac{\mathrm{d}r}{\mathrm{d}t}\frac{1}{1-r} \\
\psi &= \frac{\tilde{\psi}}{1-r} \\
\rho_0 &= \frac{\tilde{\psi}\left(0\right)}{\left(1-r(0)\right)\left(\tilde{\psi}\left(0\right)+\tilde{\mu}\left(0\right)\right)}
\end{aligned} \tag{13}
$$

The sampling rate now depends upon the removal rate:

$$\psi = \frac{\tilde{\psi}}{1-r}$$

$E$, and thus also $\rho_0$, is fully dependent on the removal rate, pulled death and pulled sampling rate.

$$\left(\tilde{\mu}-\psi r\right)\left(1-E\right)=\psi E$$

$$E\left(\psi+\tilde{\mu}-\psi r\right)=\tilde{\mu}-\psi r$$

$$E = 1-\frac{\tilde{\psi}}{\left(1-r\right)\left(\tilde{\mu}+\tilde{\psi}\right)}$$

$$\rho_0 = \frac{\tilde{\psi}\left(0\right)}{\left(1-r\left(0\right)\right)\left(\tilde{\psi}\left(0\right)+\tilde{\mu}\left(0\right)\right)}$$

We then obtain the birth rate:

$$\lambda = \frac{\tilde{\lambda}\left(1-r\right)\left(\tilde{\psi}+\tilde{\mu}\right)}{\tilde{\psi}}$$

We write the derivative of $E$ with respect to $r$ and the pulled rates.

$$
\begin{aligned}
\frac{\mathrm{d}E}{\mathrm{d}t} =& \frac{\mathrm{d}}{\mathrm{d}t}\left(1-\frac{\tilde{\psi}}{\left(1-r\right)\left(\tilde{\mu}+\tilde{\psi}\right)}\right) \\
=& \frac{\mathrm{d}\frac{\tilde{\mu}}{\tilde{\psi}}}{\mathrm{d}t}\frac{\tilde{\psi}^2}{\left(\tilde{\mu}+\tilde{\psi}\right)^2\left(1-r\right)}-\frac{\mathrm{d}r}{\mathrm{d}t}\frac{\tilde{\psi}}{\left(1-r\right)^2\left(\tilde{\mu}+\tilde{\psi}\right)}
\end{aligned}
$$

This allows us to obtain $\mu$.

$$
\begin{aligned}
\mu =& \tilde{\mu} + \lambda E - \psi r + \frac{\frac{\mathrm{d}E}{\mathrm{d}t}}{1-E} \\
\mu =& \tilde{\mu} + \tilde{\lambda}\left(\frac{\tilde{\mu} - \tilde{\psi} r - \tilde{\mu} r}{\tilde{\psi}}\right) - \psi r + \frac{\mathrm{d}\frac{\tilde{\mu}}{\tilde{\psi}}}{\mathrm{d}t}\frac{\tilde{\psi}}{\tilde{\mu}+\tilde{\psi}} - \frac{\mathrm{d}r}{\mathrm{d}t}\frac{1}{1-r} \\
\mu =& \tilde{\mu}\left(\frac{\tilde{\psi} + \tilde{\lambda} - \tilde{\lambda} r}{\tilde{\psi}}\right) - \tilde{\lambda} r - \frac{\tilde{\psi} r}{1-r} + \frac{\mathrm{d}\frac{\tilde{\mu}}{\tilde{\psi}}}{\mathrm{d}t}\frac{\tilde{\psi}}{\tilde{\mu}+\tilde{\psi}} - \frac{\mathrm{d}r}{\mathrm{d}t}\frac{1}{1-r}
\end{aligned}
$$

We can confirm that the previous solutions for non-removal sampling are special cases of these general solutions.
